# Cell type and cell signaling innovations underlying mammalian pregnancy

**DOI:** 10.1101/2024.05.01.591945

**Authors:** Daniel J. Stadtmauer, Silvia Basanta Martínez, Jamie D. Maziarz, Alison G. Cole, Gülay Dagdas, Gilbecca Rae Smith, Frank van Breukelen, Mihaela Pavličev, Günter P. Wagner

**Author notes:** These authors contributed equally (Daniel J. Stadtmauer, Silvia Basanta Martínez).

## Abstract

How fetal and maternal cell types have co-evolved to enable mammalian placentation poses a unique evolutionary puzzle. Here, we present a multi-species atlas integrating single-cell transcriptomes from six species bracketing therian mammal diversity. We find that invasive trophoblasts share a gene-expression signature across eutherians, and evidence that endocrine decidual cells evolved stepwise from an immunomodulatory cell type retained in *Tenrec* with affinity to human decidua of menstruation. We recover evolutionary patterns in ligand-receptor signaling: fetal and maternal cells show a pronounced tendency towards disambiguation, but a predicted arms race dynamic between them is limited. We reconstruct cell communication networks of extinct mammalian ancestors, finding strong integration of fetal trophoblast into maternal networks. Together, our results reveal a dynamic history of cell type and signaling evolution.

**Synopsis:** The fetal-maternal interface is one of the most intense loci of cell-cell signaling in the human body. Invasion of cells from the fetal placenta into the uterus, and the corresponding transformation of maternal tissues called decidualization, first evolved in the stem lineage of eutherian mammals(*1*, *2*). Single-cell studies of the human fetal-maternal interface have provided new insight into the cell type diversity and cell-cell interactions governing this chimeric organ(*3–5*). However, the fetal-maternal interface is also one of the most rapidly evolving, and hence most diverse, characters among mammals(*6*), and an evolutionary analysis is missing. Here, we present and compare single-cell data from the fetal-maternal interface of species bracketing key events in mammal phylogeny: a marsupial (opossum, *Monodelphis domestica*), the afrotherian *Tenrec ecaudatus,* and four Euarchontoglires - guinea pig and mouse (Rodentia) together with recent macaque and human data (primates) (*4*, *5*, *7*). We infer cell type homologies, identify a gene-expression signature of eutherian invasive trophoblast conserved over 99 million years, and discover a predecidual cell in the tenrec which suggests stepwise evolution of the decidual stromal cell. We reconstruct ancestral cell signaling networks, revealing the integration of fetal cell types into the interface. Finally, we test two long-standing theoretical predictions, the disambiguation hypothesis(*8*) and escalation hypothesis(*9*), at transcriptome-wide scale, finding divergence between fetal and maternal signaling repertoires but arms race dynamics restricted to a small subset of ligand-receptor pairs. In so doing, we trace the co-evolutionary history of cell types and their signaling across mammalian viviparity.

## Main Text

The origin of novel tissues and organs is essential for the evolution of complex multicellular organisms. However, understanding how evolution at higher levels of organization relates to evolution of cellular composition and signaling is currently limited. One of the most intense loci of cell signaling in the body is the fetal-maternal interface, where cells from the fetal placenta invade and establish communication with the maternal endometrium. Placental invasion, and the corresponding transformation of maternal tissues called decidualization, first evolved in the stem lineage of placental (eutherian) mammals(*1*, *2*). In this paper, we trace the fetal-maternal interface to its evolutionary origin, and ask a fundamental question: how does a complex novelty, composed of highly interdependent constituent parts, come to be assembled in evolution?

Answering this question demands a comparative perspective. Single-cell transcriptomic atlases of the human fetal-maternal interface have yielded insight into the cell-cell interaction networks governing this chimeric organ(*3–5*). However, the placenta is also one of the most rapidly evolving, and hence most diverse, characters among mammals. Here, we present and compare single-cell data from the mid-gestation fetal-maternal interface of five species bracketing key events in mammal phylogeny (**Figure 1a**). These include the gray short-tailed opossum, *Monodelphis domestica*, a non-deciduate marsupial, and four eutherian species: the Malagasy common tenrec *Tenrec ecaudatus*, an afrotherian mammal considered to have a “primitive” form of hemochorial (invasive) placentation (*10*, *11*), and four Euarchontoglires - the guinea pig *Cavia porcellus*, the mouse *Mus musculus*, and recently published data from two primates *Macaca fascicularis*(*7*) and *Homo sapiens*(*4*, *5*).

**Fig. 1.**
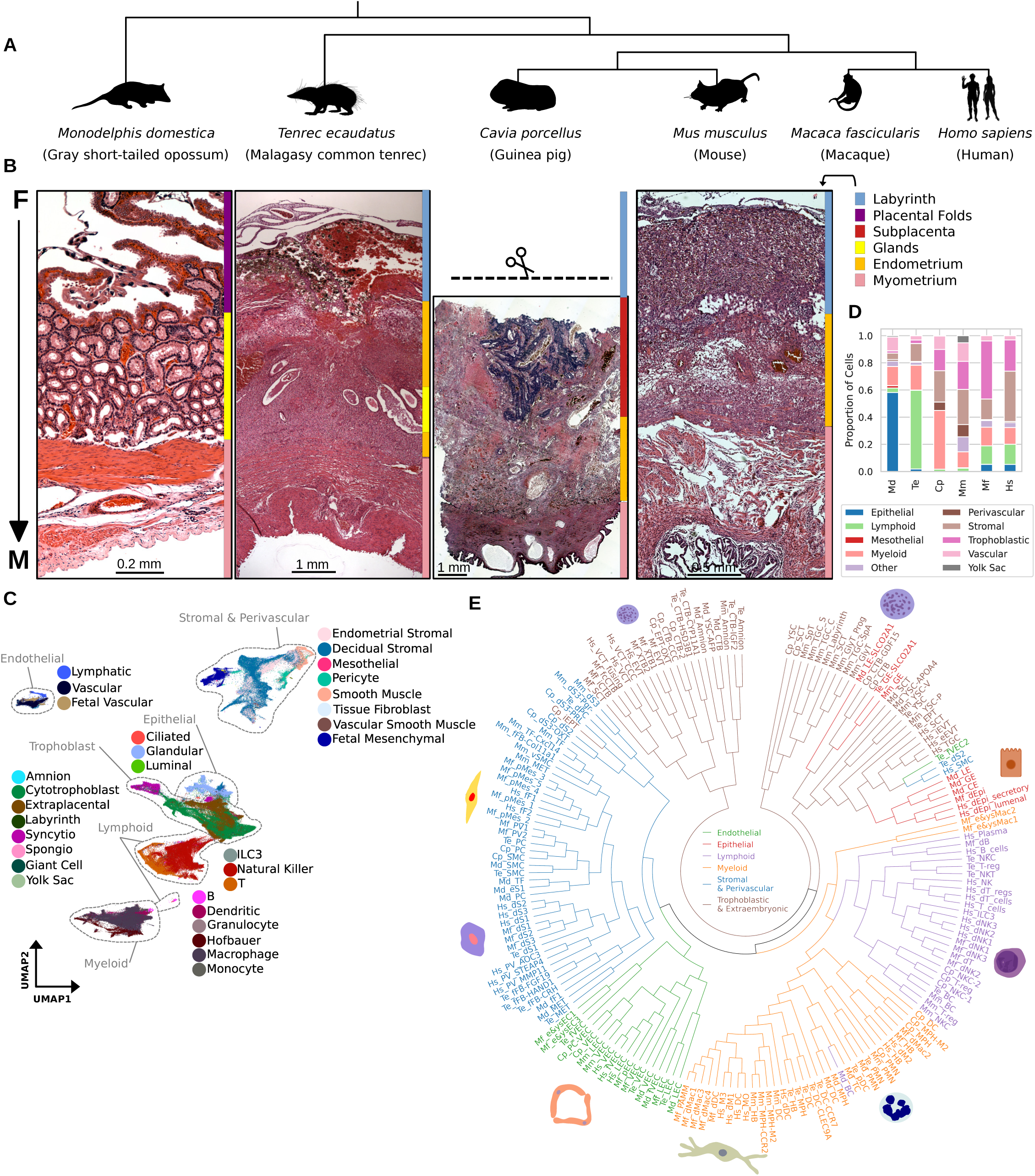
Single-cell transcriptomic atlases of five mammalian species spanning the diversification of viviparity – from left to right: *Monodelphis domestica*, *Tenrec ecaudatus, Cavia porcellus, Mus musculus, Homo sapiens*. (**A**) UMAP embeddings of single-cell transcriptomes from the species listed. (**B**) Histology of the fetal-maternal interface of the species in (a). Color bars reflect tissue regions (legend on right) from fetal (F) to maternal (M). (**C**) Unified cross-species UMAP (SAMap), colored by cell type, with prominent divisions annotated in gray. (**D**) Proportional abundances of cell types belonging to major cell type families in each sample. (**E**) A neighbor-joining phylogenetic tree showing hierarchical relationships of cell clusters recovered, colored by broad families. For a full list of cell type descriptions, see Supplementary Material. CTB: cytotrophoblast; DC: dendritic cell; dS1/dS2/dS3: decidual stromal cell; eS1: endometrial stromal fibroblast; EPT: extraplacental trophoblast; iEPT: invasive extraplacental trophoblast; EVT: extravillous trophoblast; eEVT: endovascular extravillous trophoblast; iEVT: interstitial extravillous trophoblast; fFB/fMES: fetal mesenchymal cell; GE: glandular epithelial cell; HB: Hofbauer cell; ILC: innate lymphoid cell; LE: luminal epithelial cell; LEC: lymphatic endothelial cell; MET: mesothelial cell; MO: monocyte; MPH: macrophage; NK/NKC: natural killer cell; PC/PV: pericyte; PMN: polymorphonuclear cell; SCT: syncytiotrophoblast; SMC: smooth muscle cell; SpT: spongiotrophoblast; TC: T cell; T-reg: regulatory T cell; TF: tissue fibroblast; TGC: trophoblast giant cell; TGC-SpA: spiral artery-remodeling trophoblast giant cell; GlyT: glycogen trophoblast cell; GlyT_Prog: glycogen trophoblast progenitor; TGC_S: sinusoidal trophoblast giant cell; TGC_C: canal trophoblast giant cell; VEC: vascular endothelial cell; fVEC: fetal vascular endothelial cell; YSC: yolk sac cell; YSC-P: parietal yolk sac cell; YSC-V: visceral yolk sac cell.

We establish putative cell type homologies, identifying trophoblast populations of guinea pig and tenrec which share a gene expression signature with primate extravillous trophoblast, and discover a primitive “predecidual” cell type in the tenrec which suggests a stepwise evolution of the decidual stromal cell. We reconstruct ancestral cell-cell signaling networks, revealing the integration of novel cell types into the fetal-maternal interface. Finally, we test two long-standing theoretical predictions, the disambiguation hypothesis(*8*) and escalation hypothesis(*9*), at transcriptome-wide scale. In so doing, we trace the co-evolutionary history of cell types and cell signaling across the origin and diversification of mammalian placentation.

## Results

### Therian mammals differ in placental and uterine cell type inventories

Single-cell transcriptomic data were collected from the uteroplacental interface of mammals at informative phylogenetic positions for the evolution of placentation (**Figure 1b**). Samples were collected after placental development had completed but before the physiological changes leading to the onset of parturition. Datasets were analyzed following a standardized approach (**Figure S1**). The resulting curated cell atlas (**Figure 1c**) consists of 404,118 cells and 145,153 nuclei totaling 549,271 libraries, with an average of 10,253 unique transcripts per library. Cell clusters were identified using Leiden clustering and annotated using marker gene expression and histology. To reduce over- or under-clustering, we used non-negative matrix factorization (NMF)(*12*) to identify coregulated gene modules (**Figure S2**). Cell clusters lacking uniquely distinguishable NMF gene modules or markers were merged. Detailed cell type descriptions are provided as Supplementary Material.

Tissue organization (**Figure 1b**) and cell type composition (**Figure 1d**) vary across the mammals in our study. In eutherian mammals, both a vascular interface where maternal circulation is made available to the fetus and an interface where invading trophoblast contacts maternal decidua are present. The latter was the focus of analysis due to greater potential for co-evolved cell-cell signaling. In the opossum, however, fetal-maternal contact and nutrient exchange occur in the same location, a lamellar matrix of trophoblast which coats maternal epithelium (**Figure S3a**). Uterine glands, which provide histotrophic nutrition, are expansive (**Figure 1b**): more than 50% of the cells captured in the opossum were epithelial, of which 85% were glandular (**Figure 1d**). Stromal cells such as fibroblasts are more scarce (8%) in opossum than in other species (**Figure 1b,d**). In the tenrec, guinea pig, and mouse, the vascular interface is a trophoblast labyrinth surrounding maternal blood (**Figure 1b**), whereas the human and macaque vascular interface consists of fetal villi reaching into a maternal blood sinus. In the tenrec, we identified glands containing mucus and leukocytes, suggesting continued histotrophic nutrition (**Figure S3b**), whereas in the other eutherians histotrophy has ceased by the time placentation is complete. Lymphoid cells, primarily NK-like innate cells, constituted 58% of the total cells recovered in this species (**Figure 1d**), which contrasts with the opossum, where no uterine NK cells could be identified. The guinea pig, mouse, macaque and human have similar cell type compositions (**Figure 1d**), but one notable structural difference: the guinea pig’s interface is organized into a folded cytotrophoblast structure called the subplacenta, which anchors the placenta to the maternal decidua (**Figure 1b**) and gives rise to invasive trophoblast(*13*).

Assessment of cross-species homology between cell populations was assisted by self-assembling manifold mapping (SAMap)(*14*), which integrates cells from all species into a unified manifold (**Figure 1c**) while accounting for complex gene homology. Pairwise SAMap mapping scores between cell clusters (**Table S1**) formed reciprocally linked groups (**Figure S4**). Conserved maternal cell types including smooth muscle, pericytes, leukocytes, endothelial, epithelial, and mesothelial cells showed consistent mapping across species. In contrast, trophoblast and decidual cell types displayed lower transcriptomic conservation across species, suggesting rapid cell type and gene expression evolution.

To obtain a global picture of cell type relationships preserving within-species hierarchical structure, we calculated a neighbor-joining tree of cell type transcriptomes. This yielded monophyletic or near-monophyletic groupings corresponding to major cell type families (**Figure 1e**; see **Methods**). The tree contains a hematopoietic cell clade with myeloid and lymphoid sub-groups, a clade of non-immune mesenchymal cells divided into one group consisting of fibroblasts, smooth muscle, and pericytes and the other of endothelial cells, and finally three mixed epithelial/trophoblast clades. These cell type family groupings were largely maintained when the genes used to generate the phylogeny were restricted to transcription factors (**Figure S5**), suggesting that the pattern derives from regulatory identity and not solely phenotypic convergence. Rapid innovation of trophoblast and decidual cell types made one-to-one homology assessment complicated, so we explored their evolutionary history with more detailed analyses.

### Placental cell types

Trophoblast cells are traditionally classified by topological location (e.g., extravillous), phenotype (e.g., glycogen trophoblast, syncytiotrophoblast), or ploidy (giant trophoblast), but whether these divisions correspond to cell type identities conserved across species is unknown.

Human trophoblast has been well documented by single-cell methods (**Figure 2a**)(*5*): cytotrophoblast of the placental villi (Hs_VCT) can undergo fusion, becoming syncytiotrophoblast (Hs_SCT), or invade, becoming extravillous trophoblast (EVT). EVTs can migrate interstitially (Hs_iEVT) into the decidua, or endovascularly (Hs_eEVT) to replace maternal spiral artery endothelium. Macaque trophoblast divide into the same three major cell type groupings of cytotrophoblast of placental villi (Mf_iCTB), fused/syncytial trophoblast (Mf_fcCTB, Mf_SCT), and extravillous trophoblast (Mf_EVT)(*7*). As not all mammals develop villi, we refer to other species’ migratory trophoblast cells as “extraplacental trophoblast” (EPT).

**Fig. 2.**
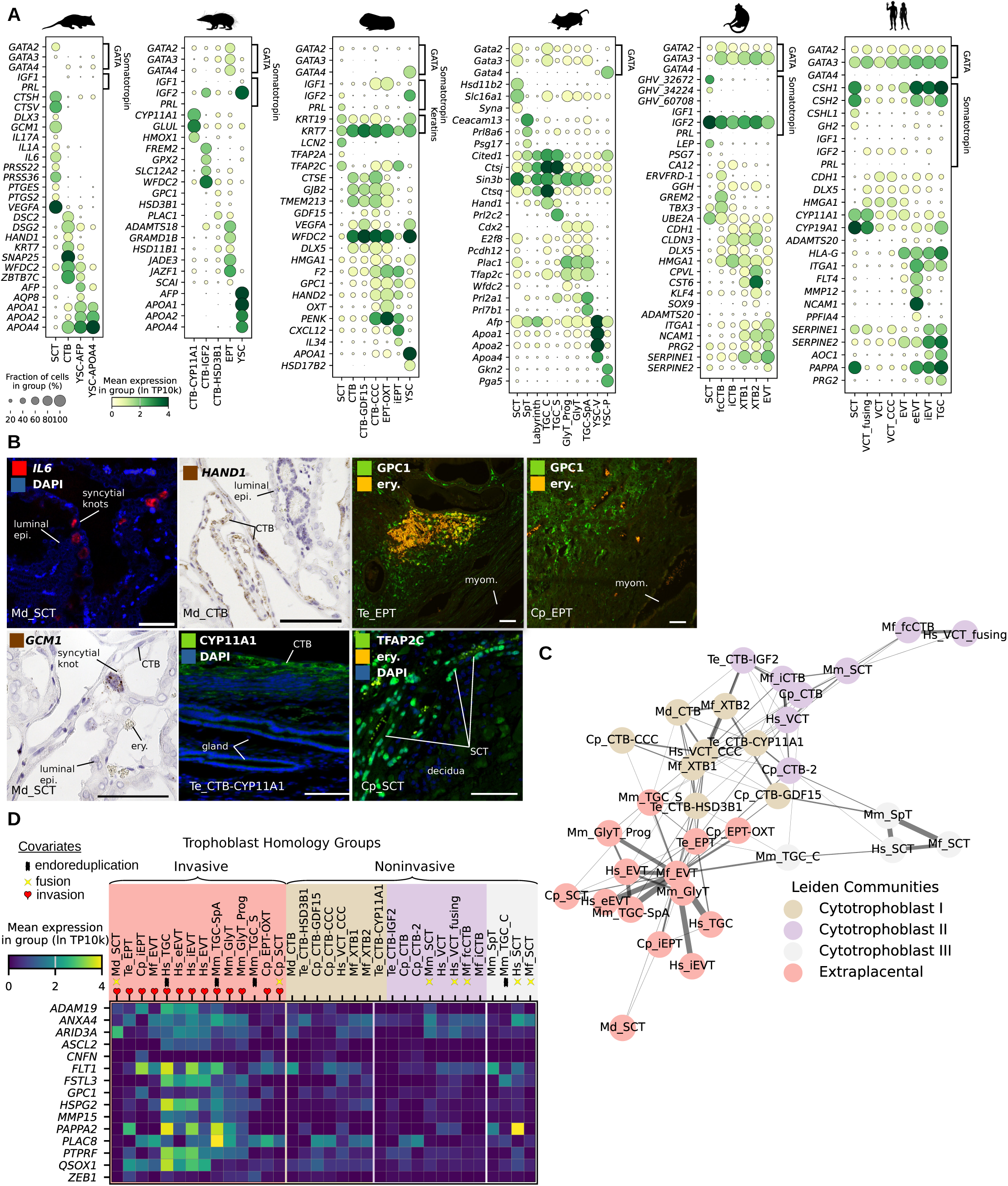
Trophoblast cell types and homology inference. (**A**) Expression of trophoblast marker genes. SCT: syncytiotrophoblast; CTB: cytotrophoblast; EPT/EVT: extraplacental/extravillous trophoblast; YSC: yolk sac cell; TGC: giant cell; CCC: cell column. (**B**) Histological localization of select cell type markers from (a). epi.: epithelium; myom.: myometrium; ery.: erythrocytes. Scale bars = 100 μm. (**C**) Network of SAMap linkages between trophoblast cells, with edge thicknesses are proportional to mapping scores. Edges with scores less than 0.1 are not shown. Nodes are colored by inferred Leiden community. (**D**) Expression of select “pan-invasive” genes in trophoblast of all species, grouped by SAMap homology group. Phenotypic covariates (endoreduplication, invasion, fusion) are annotated below cell type labels. Invasiveness groups by community, whereas fusion and endoreduplication are widespread. Md: *Monodelphis domestica*; Te: *Tenrec ecaudatus*; Cp: *Cavia porcellus*; Mm: *Mus musculus*; Mf: *Macaca fascicularis;* Hs: *Homo sapiens*.

In the opossum, two trophoblast populations were identified (**Figure 2a**). Cytotrophoblast cells (Md_CTB) express transcription factors *WFDC2* and *ZBTB7C*, desmosome components *DSC2* and *DSG2,* as well as *HAND1*, detected via in situ hybridization in mononuclear cells of the trophectoderm (**Figure 2b**). These cells also express the t-SNARE protein *SNAP25* involved in vesicle fusion(*15*), possibly related to fusion with maternal apocrine secretory bodies (**Figure S3a**). The second population consists of syncytial trophoblast, which includes giant cells of the trophectoderm, termed syncytial knots(*16*) (Md_SCT) (**Figure 2b**). Syncytial knots express *GCM1* and *DLX3,* cathepsins *CTSV* and *CTSH*, proteases *PRSS22* and *PRSS36*, high levels of the growth factor *VEGFA* and inflammatory mediators *IL1A, IL6, IL17A* and *PTGS2* as well as *PTGES*.

In the tenrec we identified four trophoblast populations (**Figure 2a**). One cluster (Te_CTB-CYP11A1) expresses *CYP11A1*, the enzyme catalyzing the first step in steroid hormone production(*17*), which we localized to the superficial-most trophoblast coating the endometrium (**Figure 2b**). These cells express heme oxidase *HMOX1*, consistent with reports in other tenrec species of catabolic breakdown of maternal hemoglobin by cytotrophoblast(*18*). Steroid hormone-producing cytotrophoblast (Te_CTB-HSD3B1) expresses the progesterone synthase *HSD3B1* and the cortisol/cortisone converting enzyme *HSD11B1*. Non-steroidogenic cytotrophoblast cells are enriched for the somatotropin *IGF2* (Te_CTB-IGF2) as well as *WFDC2*. Finally, invasive extraplacental trophoblast (Te_EPT) expresses *ADAMTS18*, *PAPPA2*, *QSOX1*, and cell invasion regulators *SCAI*(*19*) and *ZEB1*(*20*). Cells immunoreactive for the proteoglycan GPC1, a marker of invasive trophoblast(*21*) and angiogenesis(*22*), surround the deepest maternal vasculature (**Figure 2b**), suggesting deep interstitial invasion. Unlike other eutherians, no expression of *prolactin* or other growth hormone paralogs was present in any *Tenrec* placental cell type.

In the guinea pig, seven trophoblast cell clusters were distinguished. Invasion is evident as KRT7^+^ trophoblast cells are present in the decidua and surrounding maternal vasculature (**Figure S3c**). Extraplacental trophoblast (Cp_iEPT) expresses *ADAM19*, *CNFN, FLT1, QSOX1,* and *GPC1* (**Figure 2a**). GPC1^+^ cells surround maternal vasculature and reside in the myometrium (**Figure 2b**). A population of trophoblast progenitors (Cp_CTB-CCC) share expression of proliferation regulators *DLX5*(*23*) and *HMGA1* with human cell column cytotrophoblast (Hs_VCT-CCC); indeed, cell aggregations in the guinea pig subplacenta have been identified as homologs to the human cell column(*24*). Non-invasive cytotrophoblasts share the expression of the genes *CTSE, GJB2,* and *TMEM213* (**Figure 2a**). Syncytiotrophoblast (Cp_SCT) expresses three paralogs of *prolactin*, transcription factors *TFAP2A and TFAP2C*, and *LCN2*, a gene linked to invasion in human EVT(*25*). Syncytiotrophoblast “tongues”, identified by elongate cell morphology combined with nuclear TFAP2C expression (**Figure 2b; Figure S3d**), derive from subplacental cytotrophoblast and invade deeply into the decidua(*13*). Coarse-meshed syncytiotrophoblast between the subplacenta and the labyrinth and giant cells between the subplacenta and labyrinth and invading the decidua, are present histologically (**Figure S3e**) but were not captured transcriptomically due to their size.

Mouse trophoblast cells were grouped into eight populations: spongiotrophoblast, syncytiotrophoblast, labyrinth cytotrophoblast, glycogen trophoblast, and three populations of giant cells (**Figure 2a**). A subset of trophoblast are invasive, united by the expression of *Cdx2*, *Pcdh12*(*26*), *Plac1*, and *Tfap2c*. These include interstitially invasive glycogen trophoblast (Mm_GlyT), enriched in *Arid3a*, and their progenitors (Mm_GlyT_Prog), enriched for *Ascl2* and *Prdm1*(*27*). Endovascularly invasive spiral artery-remodeling giant cells (Mm_TGC-SpA) express the prolactins *Prl2a1* and *Prl7b1*. Noninvasive cells of the labyrinth, canalicular (TGC-C) and sinusoidal (TGC-S) giant cells, express *Cited1*(*28*) and *Ctsj*(*29*) (**Figure 2a**). Syncytiotrophoblast (Mm_SCT), in close proximity to maternal blood in the labyrinth, share some of these noninvasive cell markers and are also marked by *Slc16a1*, *Hsd11b2,* and the syncytin *Syna*(*30*).

### Placental cell type homology inference identifies a eutherian radiation of invasive trophoblasts

We analyzed fine-grained relationships among trophoblasts by building a network of SAMap scores and identifying reciprocally linked cell clusters using Leiden community detection (**Figure 2c**). The Leiden algorithm grouped the 35 total trophoblast cell populations into five communities which we considered putative homology groups. The resulting communities divided by degree of invasiveness, with one consisting of invasive cell types and three of noninvasive cell types (**Figure 2d**).

The grouping of invasive (extraplacental) trophoblast was driven by genes demonstrated to regulate cell invasion, including proteases *ADAM19*, *MMP15*(*31*), and *PAPPA2*(*32*), *ANXA4*(*33*)*, GPC1*(*22*), and *PLAC8*(*34*) (**Figure 2d**; **Table S1**). This group united extravillous trophoblast from human (Hs_EVT, Hs_eEVT/iEVT) and macaque (Mf_EVT) with rodent invasive cells (Mm_GlyT, Mm_TGC-SpA, Cp_iEPT) Opossum syncytial knot cells (Md_SCT) and extraplacental trophoblast from tenrec (Te_EPT) also fell into this group, suggesting deep conservation of this invasive placental gene signature across therians. Guinea pig SCT, the only syncytial cell type of the 6 species studied to have invasive properties, was also included in this homology group.

Three communities of noninvasive - primarily cytotrophoblast - cell types were also identified (**Figure 2d**). Opossum cytotrophoblast (Md_CTB) exhibited linkage to cytotrophoblast-like cells of all other species (**Figure 2d**): tenrec cytotrophoblast (Te_CTB-HSD3B1), guinea pig subplacental progenitors (Cp_CTB-CCC), and human villous cell column progenitors (Hs_VCT-CCC). The similarity of the earlier developmental stages of human and guinea pig cytotrophoblast to marsupial cytotrophoblast suggests that eutherian-specific placental cell types may have evolved from a conserved cytotrophoblast cell type.

In contrast to invasiveness, other phenotypic traits of cells, such as fusion and endoreduplication, did not consistently sort into SAMap communities (**Figure 2d**). Human giant cells (Hs_TGC), which develop from deeply migratory extravillous cells(*5*), mapped to extraplacental cells of other species rather than to giant cells of the mouse. Furthermore, placental cell types which undergo cell fusion, “syncytiotrophoblast”, did not show cross-species conservation except between human and macaque, the two species in our study with the most recent evolutionary divergence (**Figure 2c**). These formed a homology community with mouse spongiotrophoblast, driven by expression of large numbers of pregnancy-specific glycoproteins, although independent evolution of *PSG* and *CEACAM* gene families in rodents and primates(*35*) suggests that cell type homology based on expression of these genes is unlikely. While giant opossum knot cells (Md_SCT) are histologically syncytial, their gene expression differs from other therian trophoblasts by inflammatory and growth factor production. Likewise, guinea pig SCT clustered with other invasive cell types rather than with noninvasive mouse or human SCT. We conclude that, with the exception of macaque and human SCT, syncytial trophoblasts and giant trophoblast cell types most likely arose independently in major mammalian lineages.

Overall, the patterns we observe are consistent with eutherian invasive trophoblasts having descended from a single cell type with invasive potential in the therian common ancestor. This cell type was evidently retained in *Tenrec* and *Monodelphis,* which both have only one cell population belonging to invasive homology groups. This ancestral extraplacentally migratory cell likely radiated into interstitially migratory and artery-remodeling subtypes in Boreoeutherian mammals, and acquired distinct morphological phenotypes in different lineages such as endoreduplication. Conservation of a suite of invasion-enhancing genes, including matrix proteases and metastasis drivers (**Figure 2d**), suggest that an ancient potential for cellular invasion was elaborated upon during this process.

### Cross-species decidual cell diversity and a predecidual cell type in Tenrec

Decidualization is a coordinated transformation of the endometrium during pregnancy, including arterial remodeling, recruitment of tissue-specific leukocytes (NK cells), and differentiation of endometrial stromal fibroblasts into epithelioid decidual stromal cells. Decidualization is histologically observed only in mammals with hemochorial placentation(*1*). Molecular mechanisms of decidual development are poorly known outside of humans and mice, but regulatory interactions essential for decidual cell development, such as protein-protein interactions between HOXA11 and FOXO1, are shared among eutherian mammals and lacking in marsupials(*36*, *37*). Using our transcriptomic atlas, we investigated whether the decidual stromal cell is indeed a unitary cell type novel to eutherians and whether the gene-regulatory signature of decidual stromal cells is conserved across eutherian mammals.

Recent single-cell studies(*4*, *5*) suggest greater human decidual stromal cell diversity than the single prolactin-producing cell type canonically recognized in the field(*38*). We used matrix factorization to identify co-regulated gene sets within the human endometrial stroma: this identified gene expression modules corresponding to the three decidual cell populations annotated by Vento-Tormo and colleagues(*4*) (**Figure S2; Table S2**). The gene expression module of “Type I” stromal cells (Hs_dS1) includes contractility- and myofibroblast-related genes *ACTA2* and *TAGLN* and myosin light chain kinase *MYLK*. The “Type II” (Hs_dS2) gene expression module includes *IL15*, *HAND2, MEIS1, FOXO1*, and *LEFTY2.* These genes are upregulated during the initial wave of human decidualization(*39*) and commonly called “predecidual”(*40*). The “Type III” (Hs_dS3) gene expression module includes *IGFBP1* and *PRL*(*38*), as well as *WNT4* and the decidual proteoglycans *DCN* and *LUM*; we refer to these as “endocrine decidual cells”. Comprehensive sequencing of stromal cells across the human menstrual cycle found that predecidual cells generated during the secretory phase in the absence of an embryo, or spontaneous decidual cells, are *IL15^+^ PRL^-^* (*41*), similar to dS2 of pregnancy, not dS3. Thus, human *IL15^+^ PRL^+^* endocrine decidual cells are pregnancy-specific, whereas *IL15^+^ PRL^-^*predecidual cells also develop spontaneously during the proliferative phase of the menstrual cycle. The decidua of the pregnant macaque, an Old-World Monkey with similar reproductive characteristics including menstruation, shows the same three stromal cell types: *ACTA2^+^* Mf_dS1, *IL15^+^ PRL^-^* Mf_dS2, and *PRL^+^ IGFBP1^+^* Mf_dS3.

In the opossum, two stromal cell populations are present. Both express *FOXO1*, *HOXA11,* estrogen receptor *ESR1* and progesterone receptor *PGR* (**Figure 3a**). Endometrial fibroblasts (Md_eS1) are enriched for *SMOC2*, *WNT5A,* and the relaxin receptor *RXFP1*, and reside in the stroma surrounding endometrial glands(*36*). The other population (Md_TF) are enriched for *FBLN1*, *CLEC3B*, *MFAP4*, and are present in the myometrium (**Figure 3b**) and we thus consider them tissue fibroblasts. 29% of Md_eS1 cells express *HAND2* and *IL15*, markers of human predecidual cells thought to be eutherian-specific(*42*), but these cells did not group together consistently with sub-clustering. This suggests that *HAND2 and IL15* expression is a developmentally accessible gene-regulatory state even in stromal cells of the opossum, but became stabilized into a persistent cell type identity only during eutherian evolution.

**Fig. 3.**
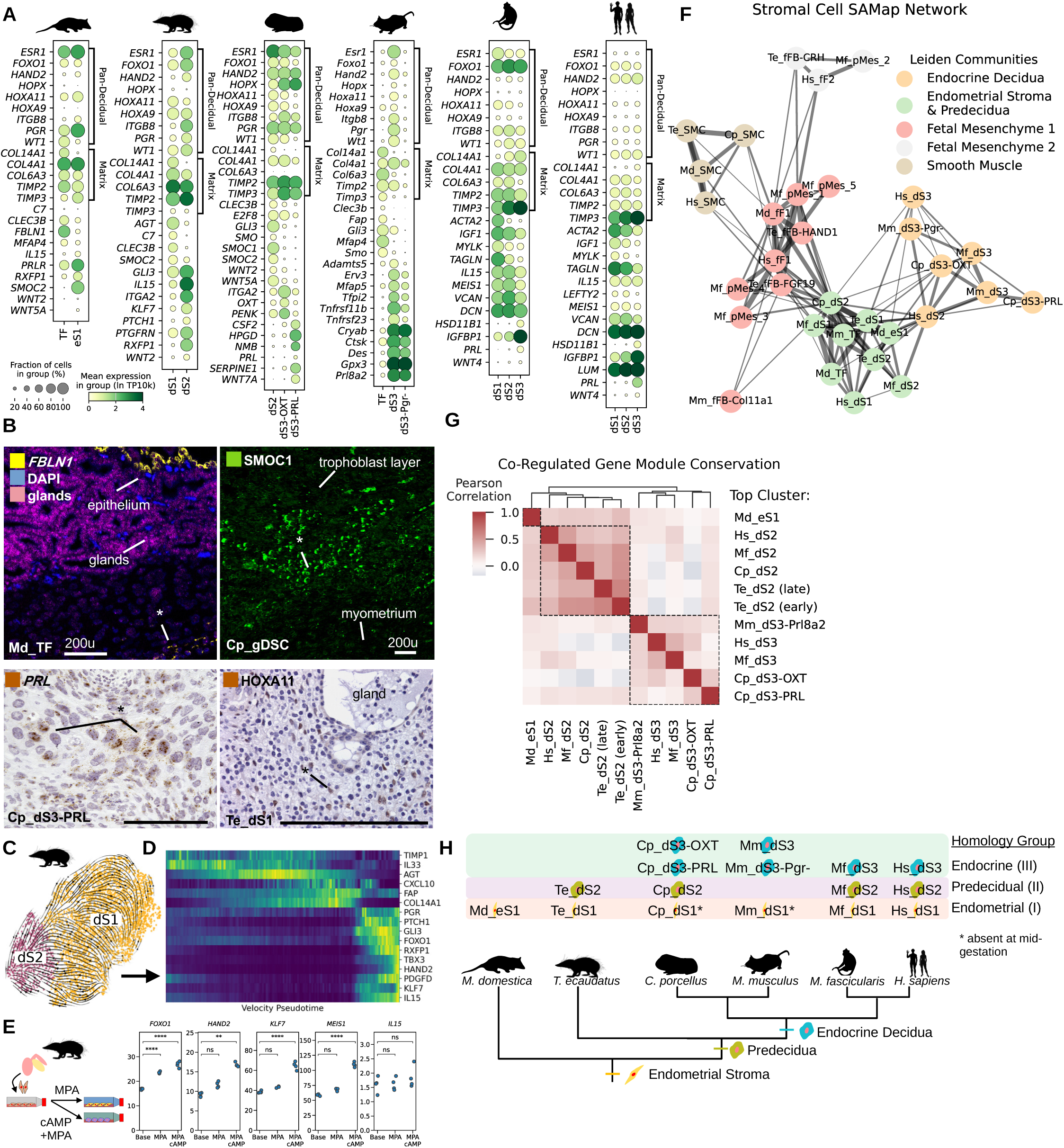
Cross-species comparison of decidual stroma. (**A**) Marker gene expression of endometrial stromal and decidual cell types. (**B**) Comparison of human midgestation decidual cells to those generated spontaneously during the secretory phase of the ovarian cycle shows a similarity of predecidua (dS2) to late luteal decidual cells. (**C**) *In situ* (italics) and immunohistochemical (upright) localization of select markers genes from (**A**); cells of interest are marked by an asterisk (*). (**D**) Force atlas embedding of tenrec endometrial stromal cells with RNA velocity vectors overlaid. Expression of select velocity-associated genes in the dS1→dS2 transition in all cells from the left plot, ordered by increasing velocity-informed pseudotime. (**E**) Expression of predecidual markers in primary tenrec uterine stromal cells cultured in vitro with deciduogenic stimuli (MPA: medroxyprogesterone acetate; cAMP) (**F**) Network of SAMap mapping scores between stromal cell types, with edge weight proportional to mapping scores, colored by community as identified by leiden clustering of the graph. Edges with scores less than 0.1 are not shown. (**G**) Correlation heatmap of NMF gene expression programs expressed in decidual stromal cells, with hierarchically-clustered dendrogram on top showing division into two families. (**H**) Tree showing inferred evolutionary relationships of decidual cell types.

In the tenrec, two major endometrial stromal cell populations are present, united by expression of *HOXA11* and *HOXA9* (**Figure 3a**). HOXA11 is localized to peri-glandular endometrial stroma (**Figure 3b**), like the opossum’s Md_eS1 cells. RNA velocity(*43*) analysis inferred a developmental trajectory from fibroblast-like (Te_dS1) to predecidual-like (Te_dS2) cells (**Figure 3c-d**). Tenrec dS1 are enriched in *CLEC3B*, *COL14A1, C7*, and *SMOC2,* whereas predecidual cells express hormone receptors *ERBB4*, *RXFP1, ESR1* and *PGR*, decidual transcription factor *KLF7*, and predecidual markers *HAND2* and *IL15*. IL15 is a chemokine for uterine NK cells^4^, with the potential to act via binding a receptor (*IL2RB+IL2RG*) expressed in tenrec uterine NK cells. Treatment of uterine stromal cells isolated from the pregnant uterus of *T. ecaudatus* with deciduogenic stimuli cAMP and MPA for 6 days elicited upregulation of these predecidual markers (**Figure 3e**), consistent with the predecidual cells observed *in vivo* representing this species’ full decidual response. The discovery of a cell population in *Tenrec* resembling *IL15^+^ HAND2^+^* human predecidual cells and lack of a clear endocrine decidual cell suggests that in this member of the basally divergent mammalian clade Afrotheria, uterine stroma persists in a state similar to the human predecidua of menstruation, rather than developing endocrine decidual cells.

The guinea pig decidual stroma contains one ECM-remodeling and two endocrinologically active populations. All three express *HOXA11*, *HAND2, ESR1, PGR,* and the rodent-specific decidual marker *HOPX*(*45*) **(Figure 3a)**. Endocrine decidual cells include a cluster enriched in oxytocin *OXT* (Cp_dS3-OXT) and another expressing a *prolactin* gene (Cp_dS3-PRL) surrounding maternal spiral arteries (**Figure 3b**). The majority (84%) of stromal cells, however, are marked by expression of *SMOC1*, *SMOC2*, and *WNT5A*. Based on their size (**Figure 3b**), expression of the cell cycle gene *E2F8* implicated in endoreduplication(*46*), and previous descriptions of giant decidual cells in necrotic areas of guinea pig decidua(*13*), we identify these as giant decidual cells, and based on SAMap affinity and lack of an endocrine profile, classify them as a Type II cell type (Cp_dS2).

The mouse day 15.5 decidua consists of a single endocrine cell type (Mm_dS3) marked by the prolactin paralog *Prl8a2*, *Pgr*, *Esr1*, and *Hand2*. This cell type also expresses 6 genes recently discovered to have maternally-biased parent-of-origin imprinting(*47*) – *Adamts5*, *Erv3*, *Mfap5*, *Tfpi2*, *Tnfrsf11b*, and *Tnfrsf23* (**Figure 3a**): these genes are tumor suppressors(*48*, *49*) and may represent mechanisms to regulate trophoblast invasion. A subpopulation of decidual cells show reduced expression of *Pgr, Esr1, Hand2,* and *Hoxa11* but maintain expression of stress-related genes, including the chaperone *Cryab*, the inflammatory enzyme *Ctsk*, and the oxidative stress gene *Gpx3*. These we term progesterone-resistant decidual cells (Mm_dS3-Pgr^-^). They have a matching gene expression profile with a *Pgr^-^* oxidative stress-enriched endocrine decidual cell population which a previous study identified as postmature decidua(*50*), and share similarities with so-called senescent *PGR^-^* endocrine decidual cells of the human which exist in balance with normal endocrine decidual cells(*51*). Cells corresponding to the predecidual cells of the tenrec and human were absent in the mouse.

### Stepwise evolution of the uterine decidua in Eutheria

While our cell type phylogeny (**Figure 1e**) identified high-level family groupings, it could not resolve one-to-one decidual cell type homologies. We investigated fine-scale decidual cell relationships using SAMap community detection and comparison of gene co-regulatory modules across species.

Two stromal cell type communities in the SAMap network were highly distinct from other mesenchymal cell types, smooth muscle cells and placental fibroblasts (**Figure 3f**). One community consists predominantly of endocrine decidual cells and the other of contractile fibroblasts and predecidual stromal cells. The endocrine or “Type III” community united the two endocrine decidual cell populations of the guinea pig (*OXT^+^* and *PRL^+^*), mouse *Pgr^+^* and *Pgr^-^* decidual cells, and human and macaque *PRL^+^ IGFBP1^+^* cells. The predecidual and non-decidual stromal cell community consisted of the opossum’s *SMOC2^+^* eS1 and *FBLN1^+^* tissue fibroblasts, tenrec dS1 and predecidual cells, guinea pig dS2, mouse tissue fibroblasts, endometrial stromal fibroblasts from macaque and human, and macaque predecidual cells. Human predecidual Hs_dS2 had strong mapping affinities to both the predecidual cells of other mammals and to endocrine decidua and were formally assigned to the same group as endocrine decidual cells. This result was robust across all iterations of community detection attempted, although we suspect that this more reflects the fuzziness of drawing hard boundaries in cells undergoing a continuous developmental process than it does an evolutionary difference between the predecidua of humans and other mammals.

We next looked for conservation of co-regulated gene modules in decidual cells of our 6-species comparison (**Figure S2**). A hierarchically-clustered correlation matrix of gene expression programs identified by NMF analysis grouped them into two families (**Figure 3g; Table S2**): Predecidual-like gene-regulatory modules from all species tended to weigh highly *MEIS1, WT1, ESR1*, *PGR,* and *EGFR*, and were active in human, macaque, tenrec, and guinea pig. Endocrine gene-regulatory modules, which had high weightings of *PRL, LUM*, *DCN*, *VIM*, *PPIB*, and *DUSP1*, were active in endocrine decidua of all eutherians (mouse, guinea pig, human and macaque), but no counterpart was identified in the opossum. Predecidual and endocrine gene expression modules showed low or negative correlation with one another (**Figure 3g**), meaning that genes expressed in the predecidual cells tended not to be expressed in mature decidual cells (**Figure S2**). We interpret this as evidence that predecidual cells and endocrine decidual cells are robustly distinct alternative gene-regulatory states.

Integrating these lines of evidence leads to the following model for decidual cell evolution (**Figure 3h**). Un-decidualized, contractile endometrial fibroblasts, or “Type I” cells, have homologs across therian mammals. These cells predominate in the non-pregnant uterus. Eutherians display decidual stromal cells falling into two states – predecidual or “Type II” decidual cells, which secrete immunomodulatory peptides, and “Type III” decidual cells, which secrete endocrine and growth-regulatory signals. Type II decidual cells likely represent a novel eutherian stromal cell type, and are evolutionarily modified from mesenchymal cells of the ancestral therian endometrium by upregulation of *HAND2* and *IL15*(*52*). In the two rodents sampled, no stromal cell type produces *IL15*; instead, decidual *IL15* is solely produced by macrophages. However, the deep phylogenetic age of the tenrec-human divergence suggests that the *IL15^+^* predecidual cell type was lost in rodents, or is restricted to early stages of pregnancy. These findings suggest that decidual stromal cell diversity expanded within crown Placentalia in two steps – establishment of the decidual stromal cell type in its initial predecidual *IL15^+^ PRL^-^* state, followed by later evolution of the endocrine (*PRL^+^*) decidual cells observed in rodents and primates.

### Co-evolutionary dynamics of cell communication

We next asked to what degree the cell-cell communication between fetal and maternal cell types shows evidence of co-evolution. We inferred signaling between cell types from complementary secreted ligand-receptor gene expression and traced the evolutionary history of these interactions within Theria.

#### Invasive trophoblast cells are integrated into the endometrial signaling network

The evolution of placentation is predicted to result in increased functional integration of fetal and maternal cells(*53*). As an approximation of functional integration, we compared the number of non-autocrine (“allocrine”) signaling interactions each cell type engages in (see **Methods**). Cells producing more allocrine ligands tend to also receive more allocrine signals themselves (Pearson’s *p* < 0.05 in all species except *Monodelphis*), and degrees of allocrine ligand production varied across major cell type families (one-way ANOVA *p* < 0.05 in all species except *Cavia*; **Figure 4a**). Stromal cells, particularly decidual stromal cells, display the greatest degrees of allocrine signaling. Invasive trophoblast (EVT/EPT, Mm_GlyT in mouse and Md_SCT in opossum) also show high degrees of allocrine signaling (**Figure 4a**). Macrophages and endothelial cells rank intermediately, whereas eutherian syncytiotrophoblast and immune effector cells (BC, TC, NKC, PMN) show the fewest allocrine interactions (**Figure 4a**). *Monodelphis GCM1^+^* trophoblast and the human Type III decidual cell have the highest Kleinberg(*54*) hub scores in their respective signaling networks, whereas lymphoid cells from all species rank among the lowest (**Figure S6a**). We interpret these findings to mean that fetal and maternal cell types that make direct contact have evolved to function as signaling hubs, highly integrated into the signaling network of the fetal-maternal interface, whereas peripheral immune cells without a stable niche do not evolve integration to the tissue microenvironment(*55*).

**Fig. 4.**
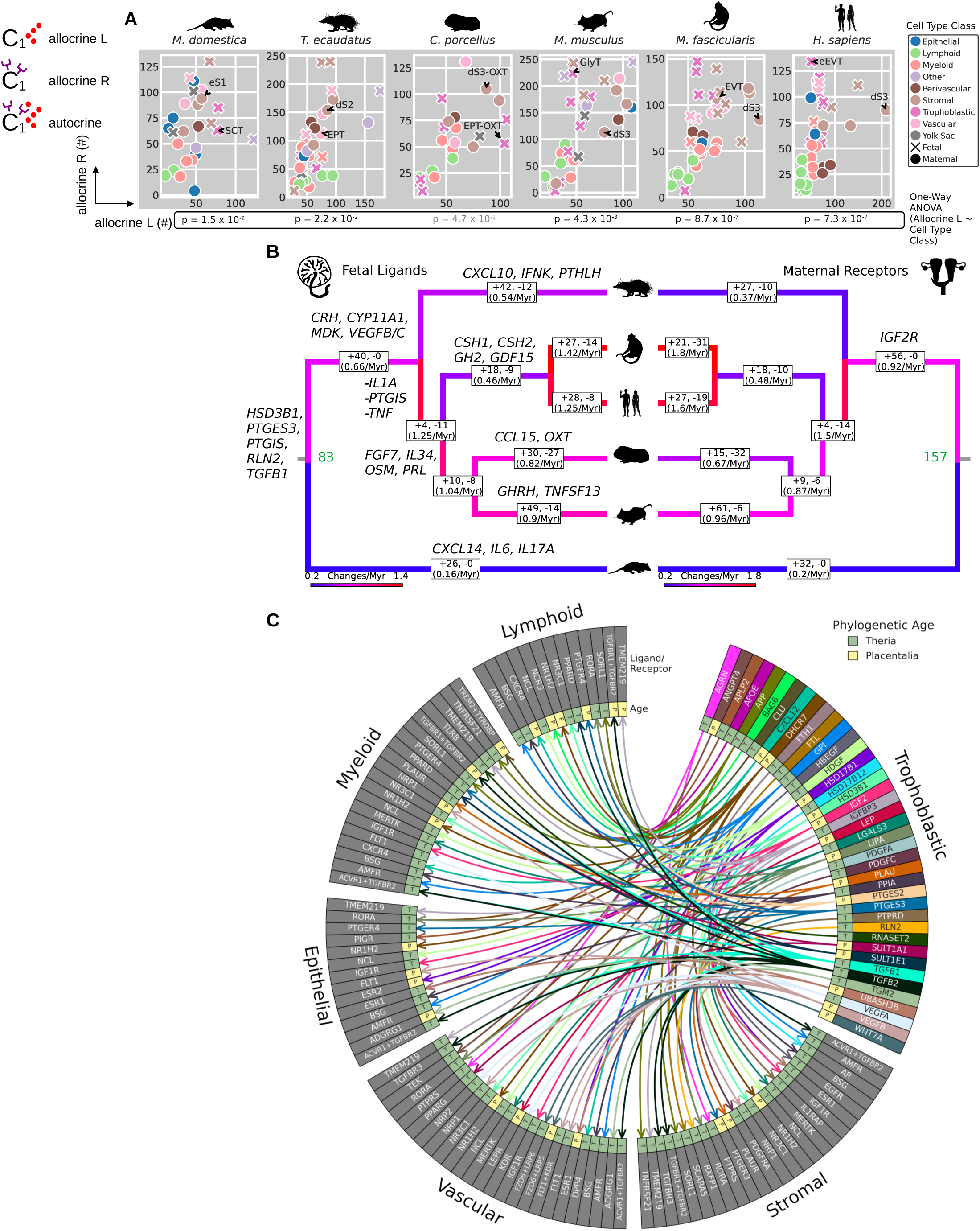
Integration of cell types into cell-cell communication networks and reconstruction of evolutionary changes. (**A**) Numbers of allocrine ligands (receptor unexpressed) and allocrine receptors (ligand unexpressed) for secreted peptide signals. Degrees of integration range from low to high and stratify by cell type class. P-values of analysis of variance relating allocrine ligand degrees to cell type class in each species are plotted below. (**B**) Co-evolution of binary expression of signals by placental cells (left tree) and cognate maternal receptors (right tree) inferred from maximum-parsimony reconstruction. Branches are colored by rate (changes/Myr) using divergence dates from (56). (**C**) Reconstructed secreted signaling interactions in the Placentalia common ancestor, colored by phylogenetic age of gene expression (eutherian novelty: P; ancestral therian: T), partitioned by cell type class and subset to only those derived from the trophoblast.

#### Reconstruction of common ancestral fetal-maternal signaling

We inferred the fetal-maternal signaling of common ancestors in our tree using maximum-parsimony ancestral state reconstruction on ligand and receptor expression of pooled fetal and maternal cell types (**Table S4**). We identify a total of 83 signals which the placenta of the first viviparous mammal likely sent to its mother, including relaxin *RLN2*, platelet-derived growth factors *PDGFA* and *PDGFC*, prostaglandin E2 via *PTGES3*, and *TGFB1* and *TGFB2* (**Figure 4b**). We traced gains and losses of signaling interactions throughout mammal evolution by comparing ancestral states. At the Placentalia node, an additional 44 signals were inferred to have been gained, including vascular growth factors *VEGFB* and *VEGFC* and the insulin-like growth factor *IGF2*. To gain more precise insight, we reconstructed full ancestral signaling networks between major cell type classes (**Table S4**). The reconstructed network of the Placental mammal common ancestor revealed that early eutherian trophoblast had already acquired a signaling potential towards maternal stromal, vascular, epithelial, and immune cells (**Figure 4c**).

Human, macaque, mouse, and guinea pig belong to the clade Euarchontoglires. Their stem lineage evolved 4 fetal-maternal signals absent in the basally branching *Tenrec,* including *CCL2* and *CCL4*, and also lost fetal expression of 11 ligands, including cytokines *IL1A* and *TNF* and neutrophil chemoattractant *CXCL2*, suggesting that inflammatory signaling by the placenta was suppressed after the divergence of tenrec and opossum. The two rodents in our sample, *Cavia porcellus* and *Mus musculus,* share an additional 10 ligands, including *FGF7*, *IL34*, *OSM* (related to *LIF*), and *prolactin*. Many fetal-maternal ligand-receptor interactions are species-specific. *Mus musculus* saw an expansion of novel *CEACAM* genes, and the midgestation guinea pig placenta is unique in *OXT* production, produced only at parturition in other species. The human lineage saw the acquisition of novel *growth hormone* paralogs, and the *Monodelphis domestica* placenta is inferred to have gained expression of additional pro-inflammatory cytokines.

We calculated rates of evolutionary change in fetal-to-maternal signaling using a time-calibrated mammalian tree(*56*) (**Figure 4b**). The greatest rates of evolutionary change were in the stem lineage leading to Euarchontoglires, with as many as 1.25 changes/Myr, many of these being losses of inflammatory signaling (e.g. *IL1A, PTGIS, CXCL2, TNF*) by fetal cell types. High rates of change were also observed in both primate lineages. In contrast, the lineage leading to *Monodelphis domestica* and the lineage leading to *Tenrec ecaudatus* show slower turnover. These findings suggest that the diversification of Euarchontoglires entailed an accelerated evolution of fetal-maternal communication compared to basally branching placentals and marsupial lineages.

#### Fetal and maternal signaling show disambiguation

A model of placental hormone evolution based on evolutionary signaling theory (*8*) predicts that fetal-maternal signaling systems will evolve towards maternal-only or fetal-only expression to reduce fetal manipulation of maternal physiology. Selection against fetal and maternal co-expression of ligands would manifest as a pattern of “disambiguation”. To quantify disambiguation, we compared the observed number of fetal-only and maternal-only expressed ligands from different signaling families (e.g. WNT, NOTCH) against a null statistical model based on random distribution of ligands to cell types regardless of origin (**Figure 5a-b**; see **Methods**). In all species, fetal and maternal co-expression of ligands was lower than expected by our null model (**Figure 5b**). The most consistently disambiguated ligand families across species included WNTs, steroids, FGFs, and various families of immunomodulatory factors (**Figure 5b; Table S3**).

**Fig. 5.**
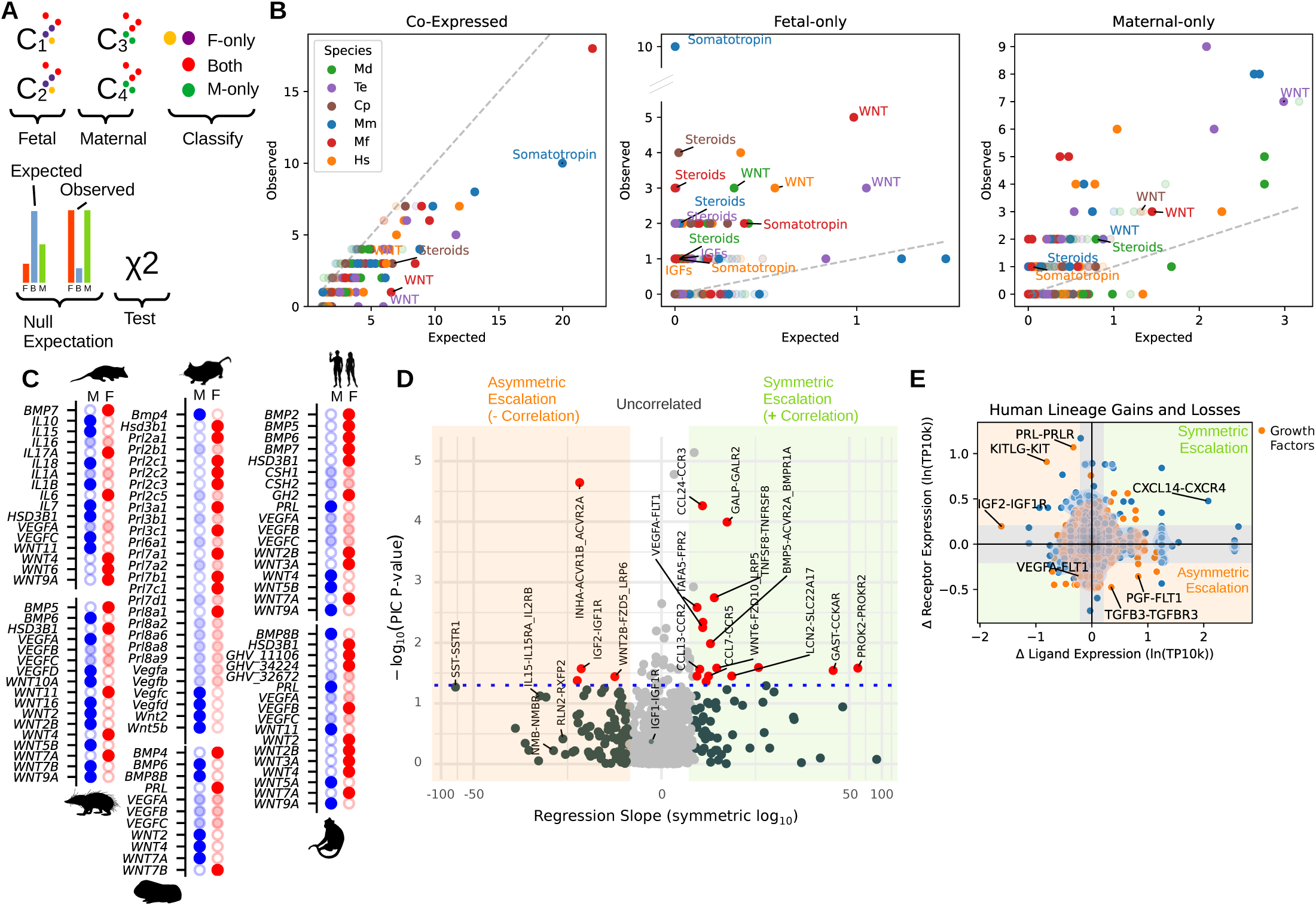
Tests of two evolutionary hypotheses for evolutionary dynamics of fetal-maternal communication, “disambiguation” (8) (**A** to **C**) and “escalation” (9) (**D** to **E**). (**A**) Design for disambiguation test. (**B**) Observed vs. expected numbers of coexpressed, fetal-only, and maternal-only ligands, by ligand family. Points with solid color have both *p* < 0.05 and Benjamini-Hochberg false discovery rate *q* < 0.05. (**C**) Disambiguation status of select ligands. Blue: maternal-only; Red: fetal-only; faint: coexpressed. (**D**) Volcano plot of ligand-receptor pairs showing significant phylogenetically-independent correlation in expression (59) on the vertical axis and correlation slope on the horizontal axis. (**E**) Inferred evolutionary changes to ligand and corresponding receptor expression in humans with respect to the human-macaque common ancestor (Catarrhini).

Gene duplication can increase disambiguation by the creation of novel placenta- or decidua-expressed paralogs. In the murine lineage, the *prolactin* gene family includes 26 members and a strongly disambiguated expression pattern with 10 fetal-specific paralogs (**Figure 5c; Table S3**). In human and macaque, duplication of the *GH1* locus has led to the origin of various placenta-specific paralogs, whereas the somatoropin *PRL* is maternal-only (**Figure 5c**). Cell type-specific growth factors involved in core tissue homeostasis, such as *VEGF*, *PDGF,* and *CSF1*, showed overwhelmingly bilateral expression (**Table S3**).

Extreme production of the angiogenic factor *VEGFA* by the opossum trophoblast can be interpreted as a nutrient-soliciting signal. The disambiguation model(*8*) predicts that in an evolutionary stable state, the mother should silence her own copy or utilize a different VEGF paralog. However, all 3 VEGF paralogs in the opossum were co-expressed. This may be explained by developmental constraints countervailing the selection for disambiguation, due to the importance of VEGF in tissue homeostasis. Alternatively, disambiguation may be achieved by maternal utilization of a splice variant of VEGFA with differential signaling properties, as documented with the *VEGF^111^* isoform in the marsupial *Sminthopsis crassicaudata*, the rat, and viviparous skinks(*57*).

#### Escalation of fetal-maternal signaling

Parent-offspring conflict over nutrient allocation during pregnancy is predicted to lead to an asymmetric evolutionary arms race(*58*) in which gains to fetal signaling strength are matched by reduced maternal responsiveness(*9*). We tested for correlated fetal ligand-maternal receptor co-evolution at the transcriptomic level by comparing reconstructed evolutionary changes in ligand and receptor expression across internal branches of the tree. We tested for a quantitative escalation pattern without complete suppression by modeling changes in expression levels as continuous traits and testing for phylogenetically-independent(*59*) associations between ligands and cognate receptor expression (**Figure 5d; Table S5**). Inferred evolutionary changes along internal branches of the tree did not show a global prevalence towards escalation regardless of whether all secreted signaling peptides are considered or just growth factors (gene ontology group GO:0008083), (**Figure S6b-c; Table S5**). Across the tree, negative ligand-receptor correlations were observed for the inhibin-activin pair INHA–ACVR1B+ACVR2A (*p* = 1.1 x 10^-3^) and the well-studied gene pair IGF2–IGF1R(*60*) (*p* = 2.7 x 10^-2^) (**Figure 5d**). Other pairs such as VEGFA–FLT1 showed positive correlation (*p* = 5.6 x 10^-3^), i.e. gains to both fetal ligand and maternal receptor expression along the same branches, possibly reflecting a cooperative trend towards greater placental vascular development in multiple eutherian lineages rather than a manipulation-desensitization dynamic.

In the branch leading from the human-macaque ancestor to humans, for which the ancestral state confidence was high due to the sampling of two rodents and two outgroups, only the growth factors *PGF* and *TGFB3* showed asymmetric escalation (**Figure 5e**), whereas *PRL*, *KITLG*, and *IGF2* were inferred to be undergoing de-escalation relative to the human-macaque ancestor. When modeling ligand and receptor expression as binary on/off traits using expression thresholding, lineages in which more fetal ligands were gained saw more gains in maternal receptors, rather than losses (**Figure 4b**). These findings suggest that only a select subset of placental-maternal signaling is subject to antagonistic gene expression evolution. That said, our analysis cannot exclude antagonistic dynamics at the post-transcriptional level, as in the case of decoy receptors(*60*) or the chemical properties of signal peptides such as their half-life(*61*).

## Discussion

The eutherian fetal-maternal interface is the product of three major evolutionary events – the origin of endometrial decidualization, the evolution of a distinct placental organ (in contrast to simple apposition of fetal and maternal tissue), and the origin of trophoblast invasion(*2*). The latter is extreme in species with deep interstitial invasion such as humans and guinea pigs. In this paper, we have compared the cell type composition and cell-cell signaling networks of species spanning these transitions revealing a complex history of cell type and cell signaling evolution.

Cross-species mapping suggests that eutherian mammals underwent an expansion in trophoblast cell type diversity. We infer that the therian ancestor, like the extant opossum, had two trophoblast cell types, one homologous to eutherian cytotrophoblast and the other specialized in signaling to the mother with weak invasive potential. Extraplacental trophoblast of tenrec, guinea pig, macaque and human and spiral artery-remodeling trophoblast in the mouse shared a common gene signature. The grouping of phylogenetically distant trophoblast transcriptomes by invasive potential suggests that the invasive/noninvasive phenotypic divide may reflect an ancient divide in trophoblast cell type identity, rather than convergent evolution.

Endometrial stromal cells fall into homology groups of different phylogenetic ages, suggesting a stepwise accrual of decidual cell diversity in eutherians. The discovery of a primitive *IL15^+^* predecidual cell in *Tenrec*, with gene expression similarity to human Type II decidual cells as well as endometrial stromal cells of other species, suggests that the predecidual cell is conserved over the 99 million year *Tenrec-Homo* divergence. Endocrine (*PRL^+^*) decidual cells likely arose later in eutherian evolution, restricted to humans and rodents in our sampling (Euarchontoglires) and lacking in *Tenrec*; investigation of decidua in more mammals from the basally-branching Afrotheria and Xenarthra clades will allow this evolution to be more thoroughly untangled.

Our findings are consistent with a model of cell type evolution playing out at two distinct levels: broad transcriptomic “cell type identities” and functional gene expression modules or “apomeres”(*62*), a division established in neuronal(*63*) and photoreceptor(*64*) evolution. In addition to novel cell types of decidual stroma and extraplacental trophoblast, fetal and maternal cells both show evidence of acquisition of functional apomeres. On the fetal side, syncytiotrophoblasts are transcriptomically dissimilar across species, and perform diverse functions - inflammatory and growth factor signaling in the opossum, invasion in the guinea pig, vascular interface in the mouse, and gonadotropin production in the human. Molecular genetic evidence suggests that trophoblast cell fusion is associated with five independent insertions of retroviral envelope genes in the opossum, tenrec, guinea pig, muroid, and primate lineages(*65–67*), a pattern congruent with our inferred lack of homology between syncytiotrophoblast beyond the human-macaque pair. Likewise, prolactin expression in endocrine decidual cells is associated with independent transposable element insertions upstream of the *prolactin* promoters in rodents and primates(*68*). These findings suggest that both trophoblast and decidual evolution have involved parallel evolution of new phenotypes, like cell fusion and hormone production, by recruitment of retroviral and transposable elements into gene regulatory networks of these cell types. Identifying which of the cell populations we identify here correspond to developmentally robust cell type identities, and the factors determining their differentiation, will require further targeted investigation.

Compelling theoretical models have long been advanced for the co-evolutionary signaling dynamics between the fetus and mother(*9*). Our test of the disambiguation hypothesis(*8*) revealed consistent and strong overrepresentation of both fetal-specific and maternal-specific ligand repertoires. This pattern was driven in part by recent gene duplication into fetal-specific paralogs(*35*). A logical next step would be to investigate genetic mechanisms, such as capture of ligand genes by fetal-specific enhancers(*69*). The escalation hypothesis predicts that increased fetal ligand production should be complemented by reduction in corresponding receptor expression by the mother. Only a select few ligand- receptor pairs in our sample consistently followed this dynamic, suggesting it does not hold for placental-uterine signaling universally; however, some of the outlier ligand-recetor pairs we identified are plausibly involved in escalation such as *IGF2*. Finally, we used ancestral state reconstruction to trace the co-evolutionary history of ligand-receptor signaling, reconstructed the cell signaling interactions of the Placental common ancestor, and found that Boreoeutherians underwent a more rapid divergence in fetal-maternal signaling than the opossum and tenrec lineages. This implies that fetal-maternal co-evolution was not solely concentrated at the origin of eutherian placentation; rather, the pace of gene-regulatory innovation in mammals has even accelerated in eutherian evolution, including substantial changes in recent primate and rodent evolution.

We acknowledge limitations to inferring ligand-receptor signaling from RNA expression. The potential for false negatives due to sequencing drop-out, incomplete genome annotation in non-model species, and protein-protein interaction divergence likely inflate the inferred number of species-specific signaling innovations. Differences in gestation length and placental physiology between our six species may limit the comparability of the chosen gestational stages for trophoblast and decidua, which are developmentally dynamic. Nevertheless, the outlines of a multistage history of cell type and cell phenotypic evolution can clearly be seen.

The fetal-maternal interface has attracted attention in recent years as a model system for single-cell genomics and ligand-mediated cell signaling at scale^3-5^. The atlas we present here demonstrates that the field has reached the point where long-standing evolutionary questions about parent-offspring co-evolution can be addressed at a resolution heretofore impossible. The molecular genetic mechanisms behind signaling innovation, integration, and disambiguation will require future investigation. As high-throughput genomics of non-traditional model species becomes more technologically accessible, expanded taxon sampling will allow the kinds of phylogenetic reconstructions we introduce here to expand in precision and scope.

The evolutionary approach we introduce for cell-cell communication holds potential for broader application. Comparative analysis may help uncover the co-evolution of cell types within other tissues, and in other inter-organismal symbioses, an avenue opened by recent advances in single-cell transcriptomics of multicellular parasites(*70*) and commensals(*71*).

## Acknowledgments

The authors thank Katie Grabek of Fauna Bio, Michael Hiller and Evgeny Leushkin for sharing the draft *Tenrec ecaudatus* genome. The study benefited from input from Caitlin McDonough-Goldstein and Jan Engelhardt. Steve Stearns and Arun Chavan provided critical comments on the manuscript. The Yale Center for Genomic Analysis (NIH 1S10OD030363- 01A1) and Yale West Campus Imaging Core supported this work.

## Author Contributions

Conceptualization: DJS, SBM, GPW, MP

Funding acquisition: GPW, MP, FvB, DJS, SBM

Experimentation: DJS, SBM, JDM, AGC, GPW

Animal Colony Maintenance: GRS, GD, JDM, DJS, SBM

Formal analysis: DJS, SBM

Software: DJS

Supervision: GPW, MP

Writing - original draft: DJS, SBM

Writing - review & editing: DJS, SBM, GPW, MP

## Data and Materials Availability

Sequencing data have been uploaded to the NCBI Gene Expression Omnibus at accession GSE274701. Human gene expression data used in this analysis were retrieved from https://www.reproductivecellatlas.org/ and macaque data from the NCBI accession GSE180637.

Data to reproduce figures and output from analyses are available in the Supplementary Materials. Analytical code for this study is available at https://gitlab.com/dnjst/fmi2024. Additional processing scripts for cell-cell communication data are archived in a python package at https://gitlab.com/dnjst/chinpy.

## Supplementary Materials

Figs. S1 to S6

Tables S1 to S8

## Materials and Methods

### Animal Husbandry

Pregnancy samples were taken at gestation points after the establishment of the definite placenta and, in the case of guinea pigs, before the luteal-placental shift in progesterone production. Time points chosen were in opossum day 13.5 of 14.5 total gestation, in tenrec day 28-29 of 56, in guinea pig day 30.5 of 62, and in mouse day 15.5 of 19.5 days gestation. These were integrated with the human atlas generated from 4-13 weeks, roughly approximate days 28-91 out of a 280-day gestation, and macaque samples generated from stages ranging from 20-140 of a 162-day gestation.

1. *M. domestica* were raised in a breeding colony at Yale University according to established technical protocols(*72*) and ethical protocols approved by the Yale University Institutional Animal Care and Use Committee (#2020-11313). Male and female animals were housed separately after 3 months of age, at which point female opossums were introduced to the male room for sexual preconditioning, and breeding was attempted after 6 months. During breeding, non-cycling female opossums were introduced to the male room for 1 day, and then subsequently swapped into the used cage of a prospective male partner for 5 days. After this period, both individuals were placed into a breeding cage and video recorded to assess the time of copulation. If multiple copulations were observed, the first was always used to calibrate 0 dpc; samples were taken one day before parturition at 13.5 dpc (n=2).
2. *T. ecaudatus* were maintained in a breeding colony at the University of Nevada, Las Vegas according to approved University of Nevada, Las Vegas Institutional Animal Care protocols. Individuals in the colony descend from a population of 40 wild-caught animals imported from Mauritius in June 2014. Animals were mated and mid-gestation was sampled for scRNA-seq between 28 and 29 days (n=3) following the first exposure of females to males.
3. *C. porcellus* (Charles River) were maintained at the University of Vienna according to Institutional Animal Care protocols, on standard chow and water ad libitum. The estrus cycle was monitored by examination of vaginal membrane opening and the animals were mated in estrus at 3-4 months of age, and video recorded to detect copulation. Midgestation samples (n=2) were collected 30.5 days post copulation.
4. *M. musculus* (C57BL/6J) were maintained at the University of Vienna on standard chow and water ad libitum according to University Institutional Animal Care protocols. Animals of age 2.5-4 months were used for the experiments. Estrus cycle was monitored by vaginal swabbing, and the females were mated when approaching estrus. Copulation was determined by the presence of a copulatory plug. The start of gestation is counted from the midnight of the night preceding the detection of copulatory plug. Midgestation samples (n=2) were collected 15.5 days post copulation.

### Single-Cell RNA Sequencing

Uteri were dissected into phosphate-buffered saline (PBS), separated from the cervix and fallopian tube, and opened longitudinally. Embryos and directly attached extraembryonic membranes were removed via severing of the umbilical cords. In the guinea pig, the labyrinth was removed prior to sample preparation. Tissue samples were kept at 4°C for approximately 30 minutes until they could be dissociated into a single-cell suspension. Due to necessity, *Tenrec* samples were shipped in RPMI medium on ice overnight and dissociated the next day.

All tissue was processed as follows, with small species-specific modifications as noted. Tissue was minced with a scalpel into ∼1 mm^3^ cubes in 2 mL of digestive solution containing 0.2 mg/mL Liberase TL (05401020001, Sigma). Tissue suspensions were heated at 37°C for 15 minutes and then passed 10 times through a 16-gauge needle attached to a 3-mL syringe. This incubation and needle passage process was repeated two more times, with the substitution of an 18-gauge needle. Finally, 2 mL of charcoal-stripped fetal bovine serum (100-199, Gemini) was added and the suspension was immediately passed through a 70-μm cell strainer then a 40-μm cell strainer. The flowthrough was pelleted by centrifugation at 500 g for 5 minutes and cells were resuspended in 1x ACK red blood cell lysis buffer (A1049201, Thermo-Fisher), incubated at room temperature for 5 minutes, centrifuged again and resuspended in PBS containing 0.04% bovine serum albumen (A9647, Sigma). At this point, cells were examined on a hemocytometer to gauge concentration and check for debris and cell death using trypan blue stain (15250061, ThermoFisher).

Mouse tissue was processed as above with the addition of Accumax (Stem Cell Technologies) to the final 0.04% bovine serum albumen solution. Guinea pig tissue was processed as with the mouse with the following modifications: a wide-bore 1 mL pipette tip was substituted for the 16-gauge needle, and 0.2 mg/mL collagenase I (17018029, Thermo- Fisher) was added to the digestion solution. As tenrec and guinea pig tissue showed higher cell death, a fractionated protocol was adopted to reduce exposure of cells to digestive enzymes. For these species, cells which had dislodged following each passage step were separated from intact tissue and immediately passed through a 70-μm cell strainer into 2 mL of charcoal-stripped fetal bovine serum (100-199, Gemini), while remaining intact tissue was allowed to continue with further digestion. Each fraction was centrifuged at 300 rpm for 5 minutes and resuspended in 500 μL ACK red blood cell lysis buffer (A1049201, Thermo-Fisher) for 5 minutes, then centrifuged at 500 rpm for 2 minutes and resuspended in 0.04% bovine serum albumen (A9647, Sigma) and kept on ice until dissociation was complete. Cells were counted on a hemocytometer and recombined in a proportional manner before sequencing.

Cells were captured using the 10X Chromium platform (3’ chemistry, version 3), libraries were generated according to manufacturer protocols (CG000315). Libraries were sequenced using an Illumina NovaSeq by the Yale Center for Genomic Analysis (*M. domestica and T. ecaudatus*) and at the Next Generation Sequencing Facility of Vienna Biocenter (*C. porcellus and M. musculus*).

### Histology and Histochemistry

Histological samples were fixed for 24 hours in 10% neutral buffered formalin, followed by dehydration to 70% Ethanol and kept at 4°C (days-weeks) or -20°C (months) before paraffin embedding. Staining with Gill 2 hematoxylin and Y alcoholic eosin was performed according to(*74*). Bright-field images for Figure 1 were taken with a Leica Thunder Imager (*Tenrec*), a Nikon Eclipse E600 (*Monodelphis*), and an EVOS M7000 (*Cavia* and *Mus*).

Immunohistochemistry was performed using antibodies recorded in **Table S6**.

For chromogenic immunohistochemistry, slides were incubated for 1 hour at 65°C, dewaxed with xylene, washed in 100% ethanol, and rehydrated in running tap water. Antigen retrieval was performed for 1 hour in a 95°C vegetable steamer containing 10 uM sodium citrate (pH 6.0). Slides were washed in phosphate-buffered saline and blocked with 0.1% mass/volume bovine serum albumen. Peroxidases were blocked by incubation in 0.03% hydrogen peroxide containing sodium azide (DAKO) in a humidification chamber for 30 minutes. Slides were incubated overnight at 4°C in primary antibody solutions, blocked, treated with horseradish peroxidase-conjugated secondary antibody for 1 hour at room temperature. Finally, slides were washed in blocking solution once more and treated with 3,3’- diaminobenzidine (DAKO K401011-2) for 5 minutes and counterstained with Gill 2 hematoxylin before brightfield imaging.

For fluorescent immunohistochemistry, the procedure had the following modifications. Peroxidase blocking and 3,3’-diaminobenzidine was not used in favor of secondary antibodies, conjugated with the fluorophore. Images were acquired using the EVOS M7000 Imaging System (Invitrogen). Fluorescent images were also captured in the red channel and superimposed to leverage the autofluorescent nature of erythrocytes to highlight the location of vasculature.

### In Situ Hybridization

Single-molecule fluorescence *in situ* hybridization was conducted on formalin-fixed paraffin-embedded sections using the RNAscope platform following the manufacturer’s protocols (ACD Bio 323100-USM for fluorescent and 322310-USM for chromogenic). Probes were designed against the equivalent transcript sequences used for RNA-seq alignment and deposited into the RNAscope database as standardized probes. Probes used are listed in **Table S7.**

Fluorescent RNAscope was performed using the MultiPlex platform (ACD Bio 323100) with Opal 520, Opal 570, Opal 620, and Opal 690 fluorophores (Akoya Biosciences). Fluorescent images were captured using a laser scanning confocal microscope (Leica Stellaris 8 Falcon). For Figure 3b, Fluorescent images were also captured at 690 nm and superimposed to leverage the autofluorescent nature of the opossum glands to highlight the location of glandular epithelia. Chromogenic RNAscope was performed using the RNAscope 2.5 HD Brown Assay (ACD Bio 322300) and imaged on an EVOS M7000 microscope.

### Single-Cell Data Analysis

Reads were obtained in FASTQ format from the respective sequencing cores. *M. fascicularis* reads were downloaded from the NCBI Gene Expression Omnibus (GSE180637), and *H. sapiens* aligned counts were downloaded from https://www.reproductivecellatlas.org.

Reads were aligned to reference genomes using the 10X Genomics CellRanger software (≥v7.0.0). *Monodelphis domestica*, *Cavia porcellus*, *Mus musculus*, and *Macaca fascicularis* were mapped to their respective Ensembl genome annotations (ASM229v1 v104, cavPor3 v104, GRCm39 v104, and Macaca_fascicularis_6.0 v112, respectively). For *Tenrec ecaudatus* a novel unpublished genome was provided by the laboratory of Michael Hiller and Fauna Bio, annotated using the TOGA pipeline(*75*).

Quality control included filtering cells with low count (less than 700 unique features for maternal cells and less than 500 unique features for fetal cells) or high mitochondrial gene expression (more than 25%), followed by doublet detection by scrublet (v0.2.3)(*76*) and doubletdetection (v4.2)(*77*), to remove clusters consisting of majority doublets. Library size normalization, log1p normalization, feature selection, and dimensionality reduction were performed using scanpy ≥ v1.9.1(*78*), with the exception of the mouse, which was annotated using equivalent functions in Seurat v4(*79*) and exported as raw counts and then reprocessed in scanpy. Principal components were adjusted to correct for batch effect across biological replicates using harmony (harmonypy, v0.0.9, r-harmony, v0.1)(*80*). UMAP embeddings were calculated and Leiden(*81*) clustering was performed to partition cells into putative cell types. Marker genes were calculated based upon differential expression (logistic regression, Wilcoxon rank-sum test, and t-test as implemented by scanpy.pp.rank_genes_groups) and used to annotate groups. Cells were annotated using a combination of these markers and active genes belonging to NMF gene expression modules (see below), and cell clusters lacking uniquely distinguishable NMF gene modules or markers were merged. Fetal cells were identified by a combination of placenta-specific marker gene expression, and non-overlap with non-pregnant uterine samples. In the case of the tenrec which was sufficiently outbred, single nucleotide polymorphism-based inference of the genome of origin using souporcell (v2.5)(*82*) (**Figure S1**). Clusters of cells which souporcell assigned to the genome of lesser abundance (putative fetal) overlapped with those annotated as fetal by differential gene expression. Spliced and unspliced transcript calling was performed using the velocyto package (v0.17.17)(*83*) and RNA velocity was modeled using scVelo (v0.30)(*43*).

Because the ground-truth database was curated from the human literature and insufficient data exist to curate ligand-receptor pairs for each species, the non-human transcriptomes were converted into “human-equivalent” transcriptomes by mapping of genes to their top human ortholog as detected by BLAST of translated peptide sequence(*14*). To maximize coverage, in cases of many:one human orthology, counts of all detected paralogs were pooled together. This transformed expression matrix was used only for cell communication analysis and cell phylogeny inference. Dimensionality reduction, cluster identification, differential expression testing, and marker gene identification and plotting were conducted on unadulterated species-relevant gene sets.

### Cell Type Phylogeny

A phylogenetic tree of cell type transcriptomes was calculated using cell type- aggregated transcriptomes from all species, with orthology resolved using the human BLAST method described above. Clusters with high annotation uncertainty, extremely low capture rate, or possible artifactual nature (’Md_LE-OAT’, ‘Te_T-gd’, ‘Md_PMN-NE’, ‘Mm_TC-NKT’, ‘Mm_NEU’, ‘Te_NEU-CHL1’, ‘Hs_Granulocytes’, ‘Mm_fMES-Myoblast’, ‘Mm_fFB-ribo’), with high erythroid signature (’Mf_Mega_progenitor’, ‘Mf_Mye_progenitor’, ‘Mf_e&ysEry2’, ‘Mf_e&ysEry3’, ‘Mf_EMP’, ‘Mf_BP’), and those originating from the embryo proper (’Mf_LP._Meso’, ‘Mf_Al’, ‘Mf_PS’, ‘Mf_PGC’, ‘Mf_Node’, ‘Mf_EPI’) were excluded from analysis. Gene expression matrices were corrected for “species signal”(*84*) by linear regression to remove the species of origin covariate from cell transcriptomes with the scanpy.pp.regress_out function. A euclidean distance matrix between cell type transcriptomes was then calculated and a neighbor-joining tree generated using the TreeMethods python package (v1.0.3; https://github.com/BradBalderson/TreeMethods). An alternative tree was also generated using the same approach with the additional subsetting of genes to only those annotated as transcription factors in Lambert et al.(*85*). Cartoons of cell types were modified from BioRender, Erkenbrack et al.(*86*), DiFrisco et al.(*87*), and original art.

### Homology Inference by SAMap

SAMap (v.1.3.4)(*14*) was used to calculate gene homology-aware similarity scores between cells of different species. BLAST (v2.14.1)(*88*) graphs were generated from the ENSEMBL proteomes of respective species, with the exception of *Tenrec ecaudatus* for which the transcriptome was used with tblastx mode. SAMap mapping tables were exported and analyzed using hierarchical clustering (seaborn, v0.13.1) as well as network analysis (networkx, v3.2.1)(*89*). For network analysis, SAMap mapping scores were used as edge weights and cell types as nodes. Communities were detected using the Leiden algorithm (leidenalg, v0.10.2)(*81*). Stability of the resulting communities was assessed by repeating community detection with a range of 10000 random initial states and constructing a co- occurrence matrix between cell types: cluster stability was defined as the mean of all co- occurrence values between members of the final community.

### Gene Expression Program Identification and Cross-Species Comparison

cNMF v1.4.1(*12*) was used to infer gene expression programs from count matrices of each species. Analysis was performed following a modified protocol from(*90*). Optimal numbers of factors (K) were chosen based upon manual examination of stability-error curves over a range of possible K values from 10-50, selecting a value immediately preceding a sudden drop in stability score (**Figure S2**).

For cross-species comparison, spectra scores from cNMF (i.e., gene loadings in NMF factors) were used. Non-human genes were mapped to their closest human orthologs as determined by BLAST score (see above), and in cases of many-to-one mapping, the highest loading score was used. Pearson correlation was calculated between programs and hierarchical clustering conducted using the clustermap function in seaborn (v0.13.1).

### In Vitro Decidualization of Primary Tenrec Stromal Cells

Mesenchymal stromal cells were isolated from the midgestation *Tenrec ecaudatus* uterus by differential attachment. Cells were expanded in T25 flasks in a growth medium consisting of 15.5 g/L Dulbecco’s modified eagle medium (30-2002, ATCC), 1.2 g/L sodium bicarbonate, 10 mL/L sodium pyruvate (11360, Thermo Fisher), and 1 mL/L ITS supplement (354350, VWR) in 10% charcoal- stripped fetal bovine serum (100-199, Gemini), and then seeded into 12-well plates (3.9 cm^2^) for 1 day. Samples were treated with either base medium (Base), 1 μM medroxyprogesterone acetate (MPA; M1629, Sigma-Aldrich), or 1 μM MPA and 0.5 mM 8-bromoadenosine 3’,5’-cyclic adenosine monophosphate (cAMP; B7880, Sigma-Aldrich) for 6 days, with replenishment on day 3 (n=4 for each condition, total n=12). Bulk RNA was isolated using a Qiagen RNeasy Micro Kit (74004), libraries were prepared by the Yale Center for Genomic Analysis and sequenced using an Illumina NovaSeq. Reads were aligned to the draft genome using kallisto (v0.45.0). Significance of differentially-expressed genes was calculated using the Wald test function of pyDESeq2 (v0.4.10)(*91*) with multiple test adjustment using the Benjamini-Hochberg method.

### Cell-Cell Communication Analysis

Putative cell interactions were inferred using expression thresholding, followed by statistical testing for significantly cell type-enriched interactions. A ground truth ligand- receptor database was built as a manually extended fork of CellPhoneDB v5.0.0(*92*) with additional curation and metadata. This modified list is archived at https://gitlab.com/dnjst/ViennaCPDB/. From these transcriptomes and the ground-truth list, interaction scores for all cell type pairs were generated using the LIANA+ (v1.2.0)(*93*) and chinpy (https://gitlab.com/dnjst/chinpy; v0.0.55) frameworks. Single-cell transcriptomes from multiple biological replicates (sample sizes reported above) were pooled before inference of cell communication was performed, following common practice(*96*). Instead of biological replicate-based testing, other statistical tests, such as CellPhoneDB(*97*) and CellChat(*98*) permutation tests, are standard; output from these tests (calculated as described above using LIANA+) are available for ligand-receptor interactions of all species as **Supplementary Data**. Likewise, the sample size of scRNA-seq studies complicates statistical testing of differential cell type abundance(*99*), so the relative abundance measures in **Figure 1d** are presented as pooled values.

For co-evolutionary integration analysis, all ligand-receptor interactions in the ground truth database were classified into “off” (ligand and receptor off), “allocrine ligand” (ligand on, receptor off), “allocrine receptor” (receptor on, ligand off), or “autocrine” (ligand and receptor on) states, based upon an expression threshold of 20% of cells in a cluster. Classification of ligands was done in this manner to measure intrinsic signaling potential for each individual cell type in a way which is not inflated by the number of clusters of other cell types identified in the tissue, in contrast to network analysis, where the splitting of a given cell type into multiple clusters inflates the number of inferred outgoing interactions for other cell types. Statistical associations between cell class and number of allocrine ligands were assessed using a one-way analysis of variance. Kleinberg hub and authority scores(*54*) were calculated for all cell types within their respective species signaling networks using the HITS algorithm as implemented in the hub_score() and authority_score() functions in igraph (v0.11.6), with edge weights set to the LIANA+ “expression products” of each inferred interaction (product of ligand and receptor log-normalized transcripts per 10,000).

For disambiguation analysis, a manually-corrected version of CellPhoneDB “classification” metadata was used to group ligands into ligand families. To preserve gene duplication events, in cases of many-to-one homology, all possible homologs to the human ground truth ligand list were included in the target species’ ligand family using Ensembl (mouse, guinea pig, opossum) or BLAST (tenrec) orthology. Expected values of disambiguated ligands were calculated as described in Statistical Analysis.

For discrete ancestral state reconstruction, interactions were scored in a boolean on/off manner based on a nonzero expression threshold of 10% of cells in a cluster for all subunits of both ligand and receptor. Maximum parsimony was calculated using the castor package (v1.8.0)(*94*). Edge changes were calculated for all internal branches of the 5-species tree by identifying genes considered “on” in descendant nodes but inferred “off” in the immediate common ancestor node, or vice versa.

For continuous ancestral state reconstruction, restricted maximum likelihood via the ape package (v5.8)(*95*) was used to estimate ancestral states. Expression values for continuous analysis were in natural log-transformed transcripts per 10,000 scale (scanpy.pp.log1p). To reduce artifacts, only genes expressed in at least 4 of 5 species were subjected to continuous ancestral state reconstruction (i.e., one loss/absence of a gene was tolerated). Changes in continuous expression values for ligands and receptors along internal branches of the 5-species tree were calculated by subtracting descendant node expression values from inferred immediate ancestral node values.

For escalation analysis, phylogenetic independent contrasts(*59*) (PIC) were calculated for ligand and corresponding receptor expression values (scaled as above) using the pic() function in ape (v5.8)(*95*). Linear regression was performed on the PIC values using the model ligand PIC ∼ receptor PIC + 0. That is, with ligand PIC as the independent variable and receptor PIC as the dependent variable, with the intercept forced to zero (assuming that when there is no change in ligand expression, there is no change expected in receptor expression). P-values reported as “PIC P-value” result from a two-sided t-test on the estimated slope coefficient of linear regression of ligand versus receptor PICs. Linear regression was also performed on observed expression values to return a regression slope.

### Statistical Analysis of “Disambiguation”

Disambiguation of ligands was compared to a null model in which the probability of randomly redistribution of ligands to cell types leading to exclusively maternal or exclusively fetal expression is assessed. First, all secreted peptide and small molecule-mediated ligand- receptor interactions were classified into “fetal-only”, “maternal-only”, or “both” expression groups based on the threshold value of 20% or more of the cells sequenced expressing the ligand in. Ligands were grouped into ligand families using an additionally curated version of the CellPhoneDB (v5.0.0) “classification” metadata. For each ligand family, the per-cell type probability of a given ligand being expressed was calculated according to the equation:

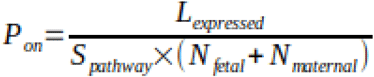

where L_expressed_ is the observed number of “on” calls within the ligand family across fetal and maternal cell types, N_fetal_ is the number of fetal cell types, N_maternal_ the number of maternal cell types, and S_pathway_ the number of ligands in the family.

The null probabilities of fetal-only and maternal-only expression for a ligand, respectively, were given by:

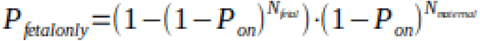

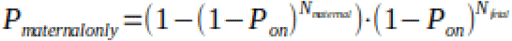

The expected fetal-only and maternal-only ligands in each family were obtained by multiplying P_fetalonly_ and P_maternalonly_ by the number of ligands in the family.

Finally, a one-way chi-squared goodness of fit test for identity of observed and expected counts of the 3 scoring categories (coexpressed, fetal-only, and maternal-only) was used to obtain chi-squared and p-values, with individual observations corresponding to ligands within a family. The Benjamini-Hochberg correction was used (statsmodels.multitest.multipletests v0.14.1) with an alpha of 0.05 to obtain false discovery rate q-values. For plotting, ligand families were considered significant if p ≤ 0.05 and q ≤ 0.05. Bonferroni-corrected p-values, obtained by multiplying *p* by the number of ligands in a family, are also reported in **Table S3**.

---

**Fig. S1.**
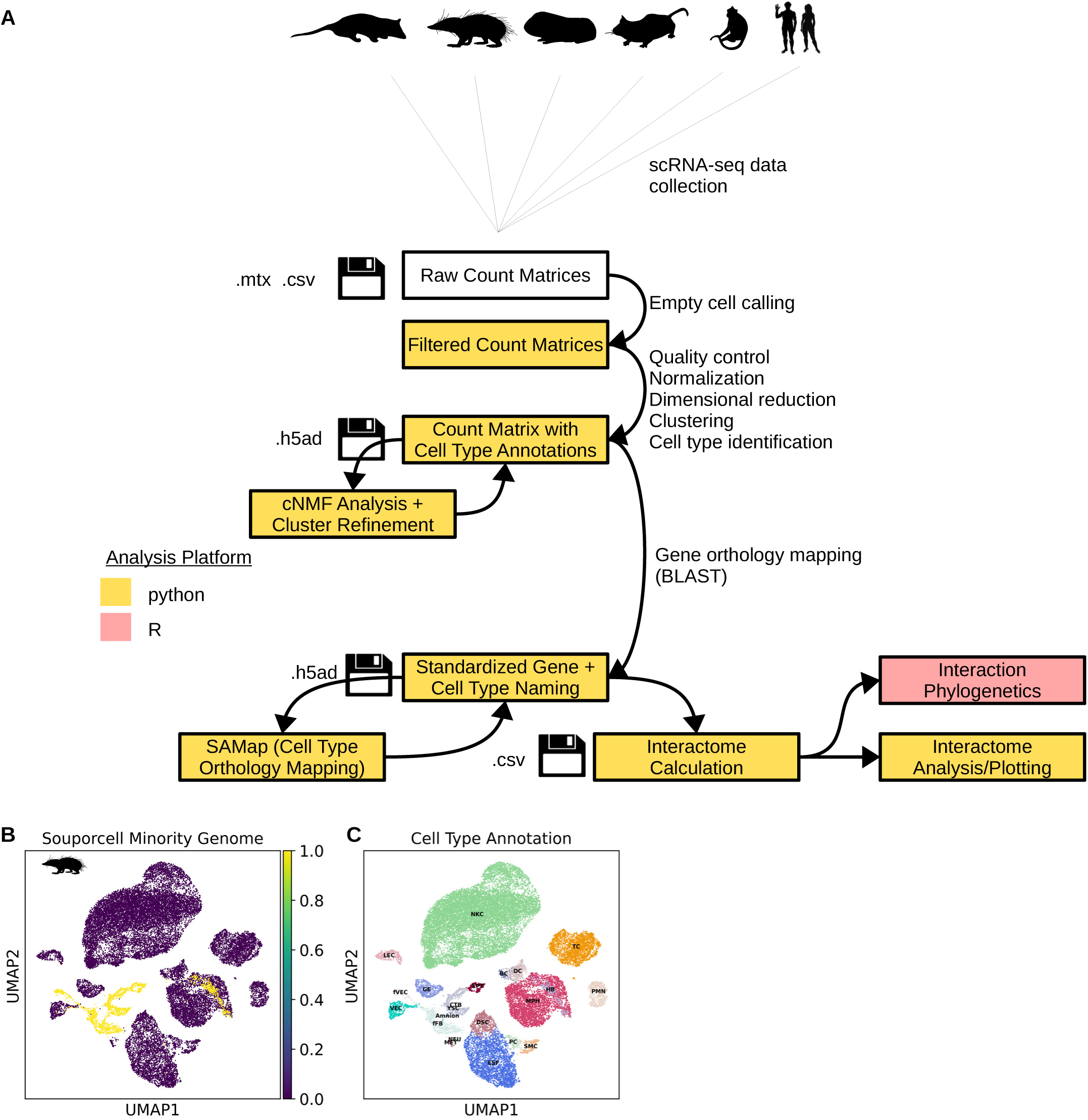
Single cell RNA-seq analysis. (**A**) Flow chart of analytical design. (**B** and **C**) Souporcell calls for *Tenrec ecaudatus* (k=2 genomes). UMAP embeddings of all cells are shown, colored by cell subtype on the left, and by whether or not the cell was assigned to the less frequent genome (assumed fetal) by souporcell on the right.

**Fig. S2.**
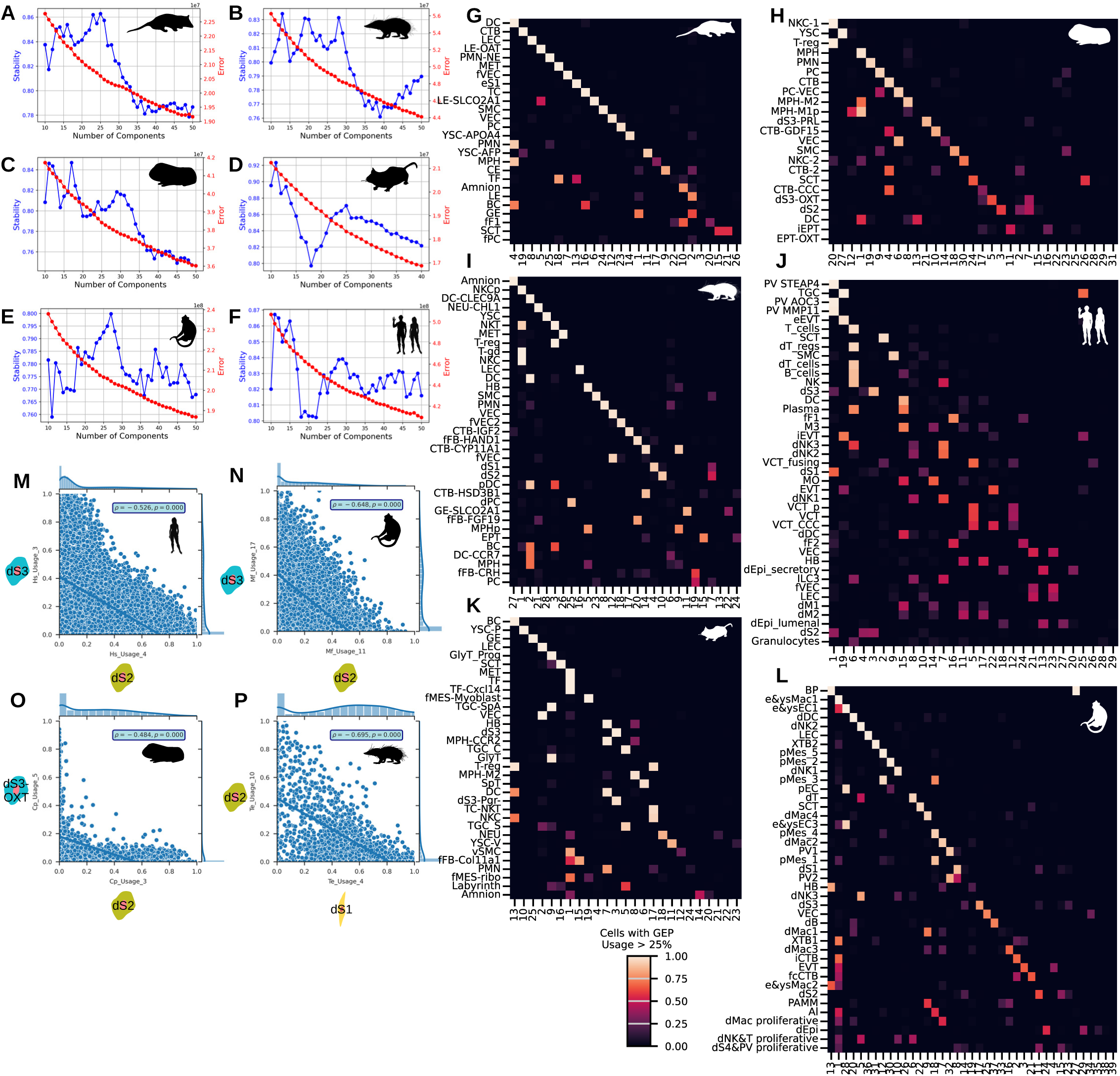
NMF analysis. (**A** to **F**) Optimal numbers of gene expression programs (NMF factors) were determined by identifying local stability values preceding a sudden drop. *Monodelphis domestica*: K=26; *Tenrec ecaudatus*: K=28; *Cavia porcellus*: K=31; *Mus musculus*: K=25*. Homo sapiens:* K=29. *Macaca fascicularis:* K=39. (**G** to **L**) Confusion matrices showing the mean activity score (ranging from 0 to 1) of cells in each cluster, taken from all cells with scores greater than 0.25. (**M** to **P**) Representative “Type II” and “Type III” decidual gene expression program activity scores across individual stromal cells of human (**M**), macaque (**N**), and guinea pig (**O**) show a diversity from exclusively Type II or Type III to a mixture of both programs. An equivalent comparison of Type I and II in the tenrec is depicted in (**P**).

**Fig. S3.**
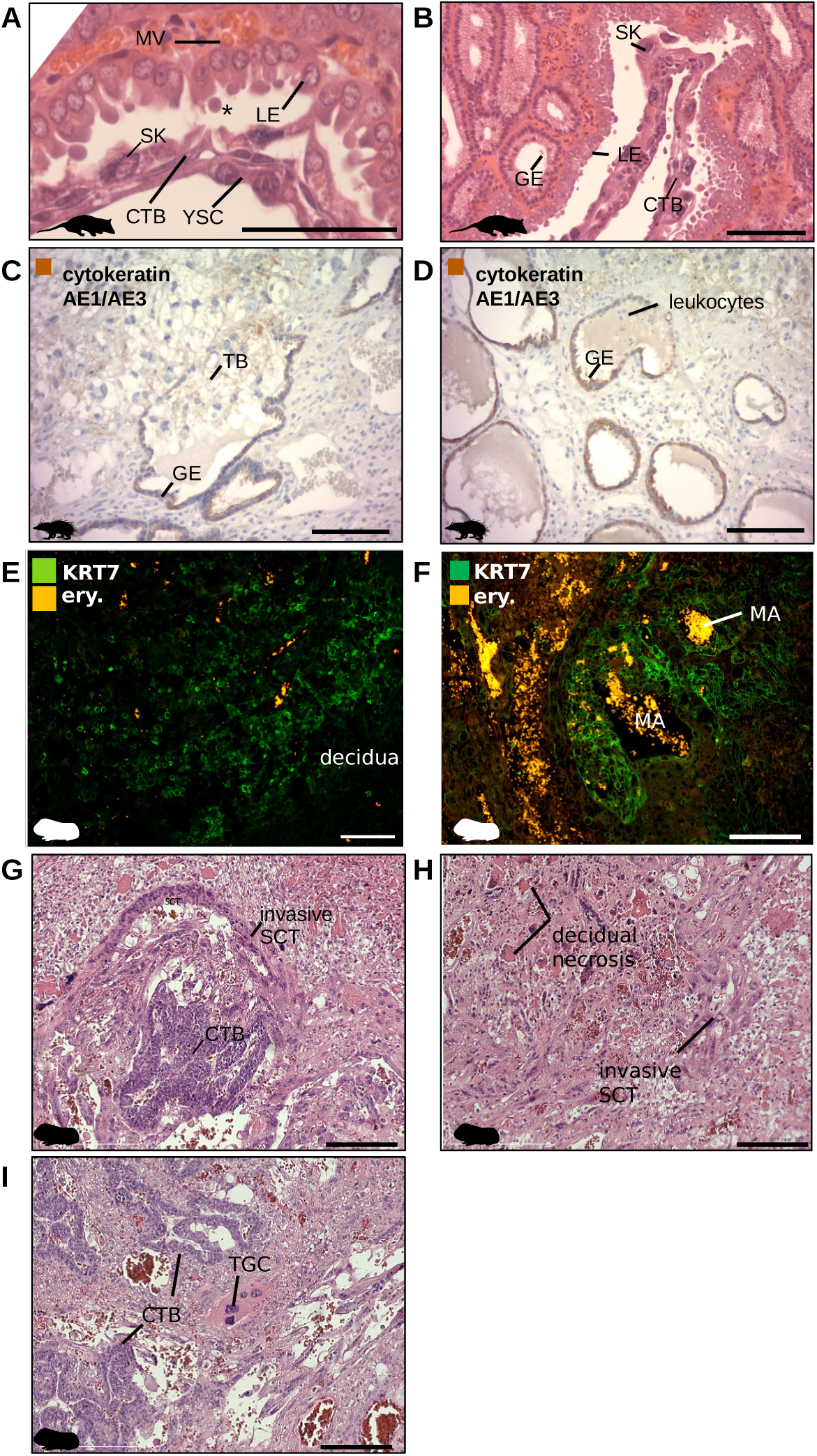
Additional histological identification of placental cell types. (**A** and **B**) The opossum fetal-maternal interface, showing 3 layers of yolk sac endoderm (YSC), cytotrophoblast (CTB), and syncytial knots (SK), in apposition with maternal luminal epithelium (LE) which produces secretory bodies (*); maternal glandular epithelium (GE) lies closely underneath the lumen. (**C** and **D**) The tenrec invasion front, where trophoblast invades into mucus-filled glands, eroding the glandular epithelium (GE) immunoreactive for pan-cytokeratin stain (CK AE1/AE3). (**E** and **F**) KRT7^+^ trophoblast in the guinea pig surrounding decidual stroma (left) and maternal arteries (MA; right). (**G** and **H**) Invasive syncytiotrophoblast (SCT) in the guinea pig emanates from subplacental cytotrophoblast (left) and extends into stroma where necrotic patches develop (right). (**I**) Trophoblast giant cell observed in the subplacenta of the guinea pig (proximate to subplacental CTB), but these cells were not captured by droplet-based sequencing. Scale bars: 200 μm.

**Fig. S4.**
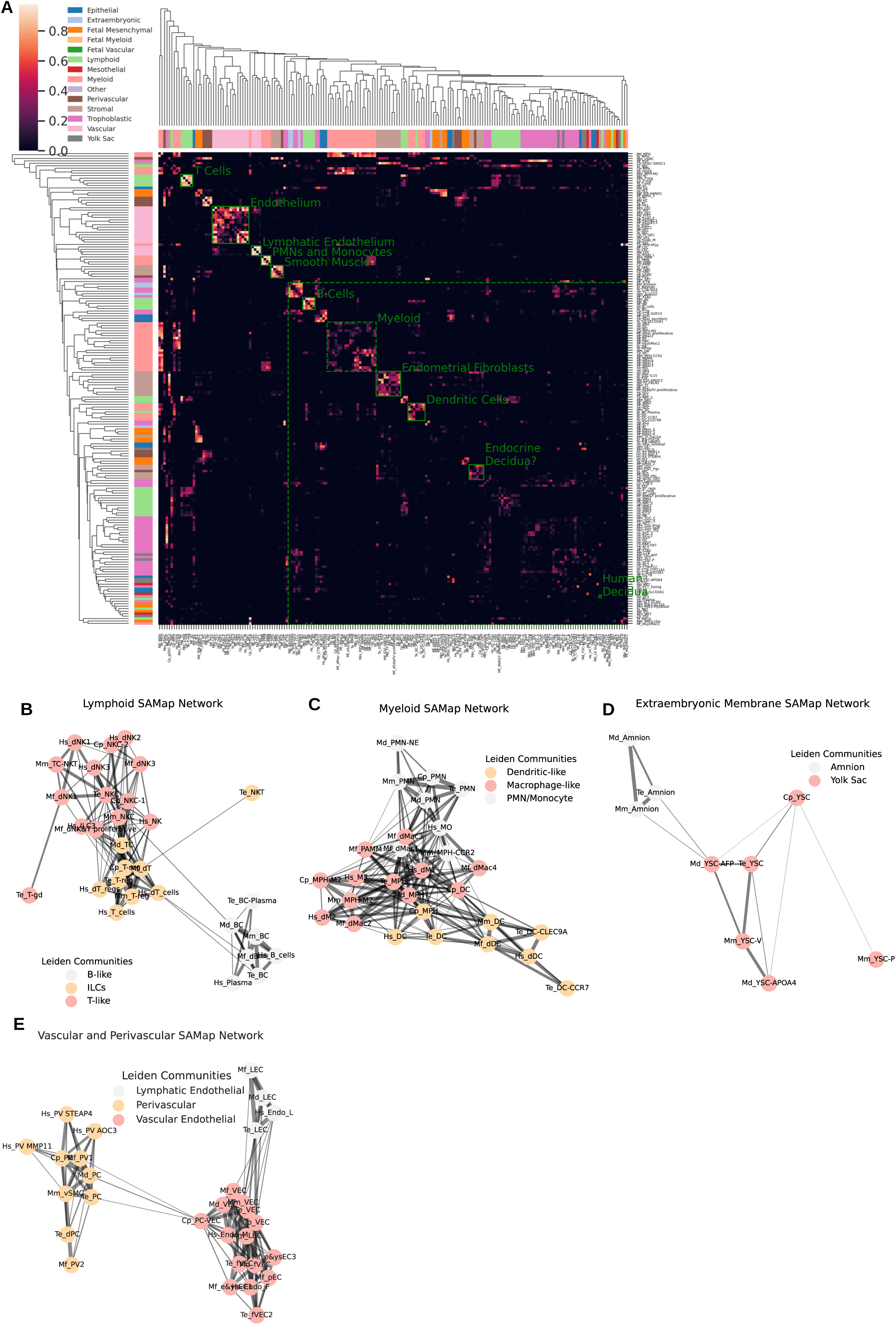
Cell type cross-species mapping scores (SAMap). (**A**) Clustered heatmap of pairwise cell type mapping scores (0–1), with selected 1:1 orthogroups annotated in green. (**B** to **E**) Networks subset from the full SAMap mapping table of (**A**) for specific cell type families, colored by Leiden community after community detection was run on the subnetwork. Nodes are cell clusters, and edge weights are equal to score magnitude, with scores less than 0.10 not shown.

**Fig. S5.**
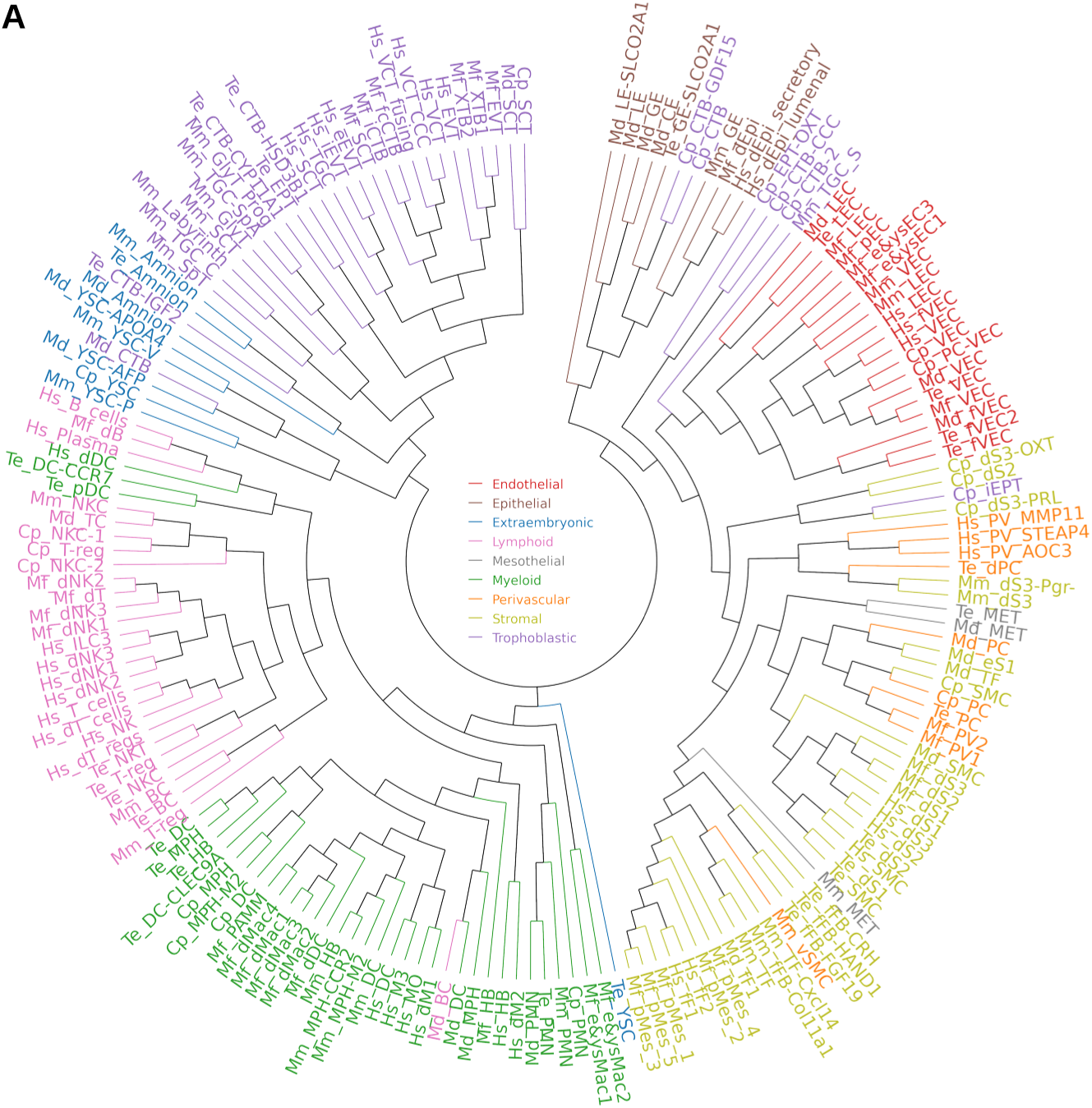
Cell type dendrogram of the cell types studied. (**A**) Neighbor-joining tree of transcription factor-only transcriptomes from all species studied.

**Fig. S6.**
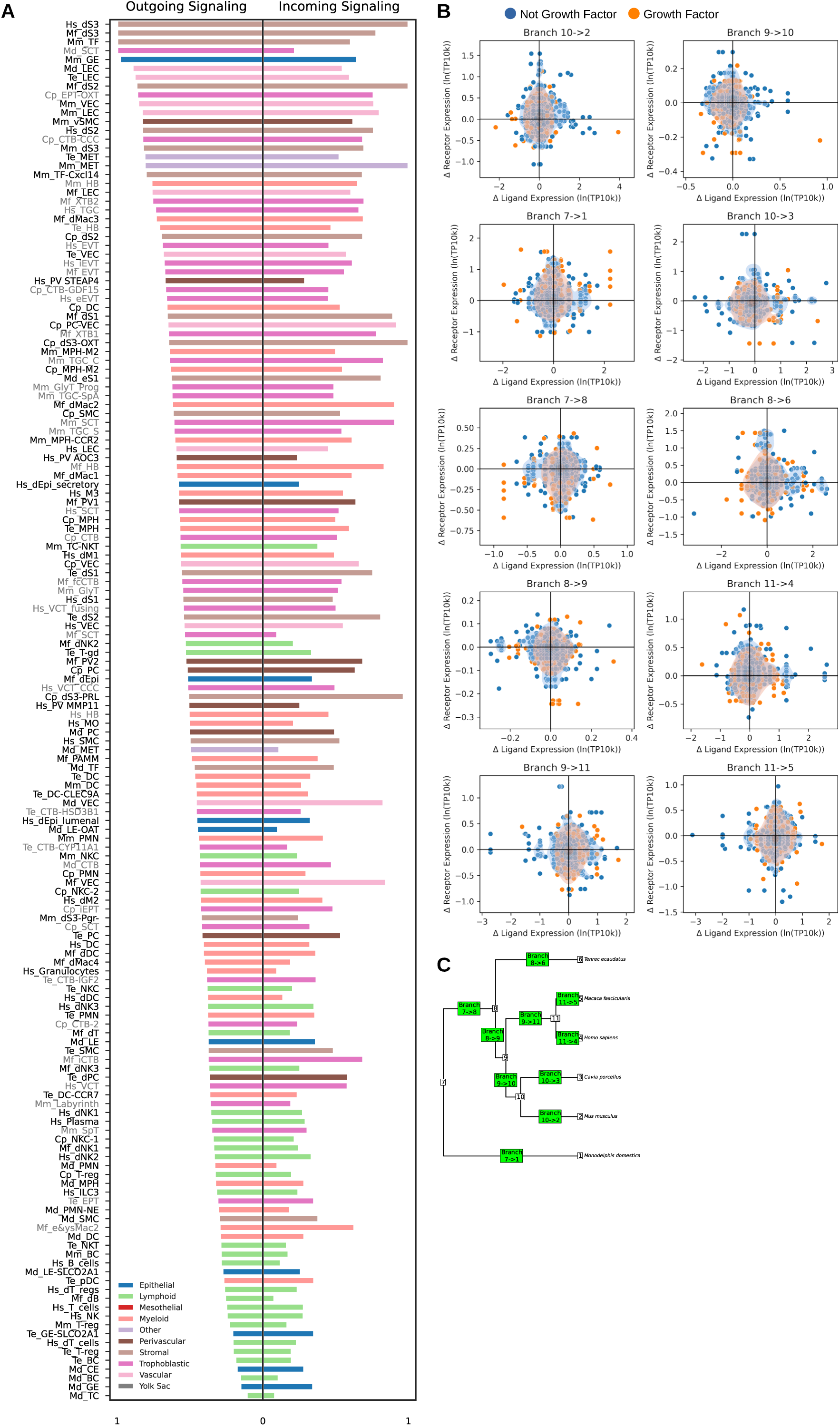
Extended cell-cell communication analysis. (**A**) Graph statistics of all cell types within their respective species’ signaling networks. Kleinberg hub scores (a measure of outgoing signaling) are shown on the left and Kleinberg authority scores (a measure of incoming signaling) on the right. Graphs are weighted based on the product of expression values of ligand and receptor. (**B**) Inferred changes in inferred expression magnitude of placental ligands and cognate maternal receptors of secreted signals along internal branches of the phylogeny. Asymmetric escalation and de-escalation events fall into the lower-right and upper-left quadrants, respectively, and uncorrelated changes cluster around the center. (**C**) Phylogenetic tree of species sampled with numbering of nodes and branches labeled as in (**A**).

